# Alternative splicing coupled to nonsense-mediated decay coordinates downregulation of non-neuronal genes in developing neurons

**DOI:** 10.1101/2023.09.04.556212

**Authors:** Anna Zhuravskaya, Karen Yap, Fursham Hamid, Eugene V. Makeyev

## Abstract

The functional coupling between alternative pre-mRNA splicing (AS) and the mRNA quality control mechanism called nonsense-mediated decay (NMD) can modulate transcript abundance. Previous studies have identified several examples of such a regulation in developing neurons. However, the systems-level effects of AS-NMD in this context are poorly understood. We developed an R package, factR2, which offers a comprehensive suite of AS-NMD analysis functions. Using this tool, we conducted a longitudinal analysis of gene expression in pluripotent stem cells undergoing induced neuronal differentiation. Our analysis uncovered hundreds of AS-NMD events with significant potential to regulate gene expression. Notably, this regulation was significantly overrepresented in specific functional groups of developmentally downregulated genes. Particularly strong association with gene downregulation was detected for alternative cassette exons stimulating NMD (NS-CEs) upon their inclusion into mature mRNA. By combining bioinformatics analyses with CRISPR/Cas9 genome editing and other experimental approaches we show that NS-CEs regulated by the RNA-binding protein PTBP1 dampen the expression of their genes in developing neurons. We also provide evidence that the NS-CE activity is temporally coordinated with NMD-independent gene repression mechanisms. Our study provides an accessible workflow for the discovery and prioritization of AS-NMD targets. It further argues that the AS-NMD pathway plays a widespread role in developing neurons by facilitating the downregulation of functionally related non-neuronal genes.

## Background

Many mammalian transcripts can undergo alternative splicing (AS), incorporating distinct exons and, occasionally, introns into mature mRNAs depending on the circumstances [1, 2]. AS plays a crucial role in the production of tissue- and cell type-specific protein isoforms [3, 4] with numerous functionally important and evolutionarily conserved examples of this regulation identified in the nervous system [5–8].

Besides diversifying the proteome, AS can regulate gene expression by changing the ratio between productively spliced mRNAs that encode functional proteins and unproductively spliced isoforms eliminated by RNA quality control mechanisms [9–11]. mRNAs containing premature translation termination codons >50-55 nucleotides (nt) upstream of an exon-exon splice junction typically undergo nonsense-mediated decay (NMD) in the cytoplasm following a pioneer round of translation [12–14]. NMD is well known for its role in destabilizing aberrant transcripts produced because of splicing errors or mutations in *cis*-elements or *trans*-acting factors involved in splicing regulation [15, 16].

Additionally, the scheduled production of alternatively spliced transcripts sensitive to NMD (AS-NMD) can facilitate the normal control of gene expression. For instance, several RNA-binding proteins use AS-NMD to auto- or cross-regulate their expression levels [17–21]. These post-transcriptional circuitries often rely on two types of alternative cassette exons (CEs) that trigger NMD when included into or skipped from mature mRNAs. We will refer to the former group as NMD-stimulating (NS-CEs; also known as “poison exons”), and the latter, NMD-repressing (NR-CEs).

AS-NMD is also known to contribute to the gene expression dynamics in development. A striking example of this regulation is provided by the large-scale upregulation of retained introns (RIs) in differentiating granulocytes and possibly other cell types, triggering NMD-mediated downregulation of functionally related genes [22–25]. Notably, bioinformatics analyses of tissue- and cell type-specific transcriptomes have identified NMD-regulating CEs in diverse groups of genes [26, 27]. Yet, it is currently unknown whether specific groups of CEs, or other exonic events such as alternative donors and acceptors (ADs and AAs), might coordinate gene expression dynamics on a scale similar to that observed for RIs.

Several lines of evidence point to the possibility of such systems-level regulation in developing neurons. For example, the RNA-binding protein PTBP1 controls NMD-repressing CEs in transcripts encoding the neuron-enriched PTBP1 paralog PTBP2 and the post-synaptic proteins GABBR1 and DLG4/PSD-95 [28–31]. These exons are predominantly skipped in embryonic stem cells (ESCs) and neural progenitor cells (NPCs) – where PTBP1 is abundant – giving rise to NMD-sensitive splice forms. The downregulation of PTBP1 in developing neurons by the neuron-enriched microRNA miR-124 facilitates the CE inclusion, thus protecting Ptbp2, Gabbr1 and Dlg4 mRNAs from NMD [29]. PTBP1 also represses NMD-stimulating CEs and ADs in several non-neuronal genes including *Bak1*, *Flna*, and *Hps1*, limiting their expression to ESCs and NPCs [32–34]. The decline in PTBP1 levels in neurons promotes the inclusion of these events, rendering the mature mRNAs sensitive to NMD.

Mutations in key NMD factors are known to lead to neurological and psychiatric defects [35–38]. NMD is also known to control neural cell identity, fine-tune the expression of important axonal and dendritic components, and regulate the excitation/inhibition balance in mature neurons [39–43]. It is tempting to speculate that these diverse functions involve coordinated regulation of different groups of AS-NMD events in a differentiation stage-specific manner.

However, understanding the developmental functions of AS-NMD at a systems level is a challenging task due to the lack of straightforward approaches for shortlisting biologically meaningful events. Moreover, NMD targets are inherently unstable and tend to be underrepresented in reference transcriptome annotations. The quantitative aspect of AS-NMD is also poorly understood. In most cases, it remains unclear whether this pathway drives developmental changes in gene expression or simply fine-tunes the abundance of productively spliced transcripts.

To systematically identify AS-NMD events with a strong potential to regulate genes in developing neurons, we developed a user-friendly computer package factR2 that annotates custom transcriptomes and prioritizes targets with significant correlation between NMD-protective splicing patterns and gene expression levels. By combining factR2 with longitudinal analysis of neuronal differentiation and acute inhibition of the NMD pathway, we uncovered a widespread role of NMD-stimulating CEs in downregulating non-neuronal genes in developing neurons. We validate our findings and demonstrate their generality using ex vivo neural cultures, subcellular fractionation, minigene and CRISPR-Cas9 experiments, and single-cell data analyses.

## Results

### An inducible system for longitudinal analysis of AS-NMD in developing neurons

We began by establishing a single-dish protocol for neuronal differentiation in vitro that avoids cell dissociation and replating steps (Fig. 1A). Inducible expression of the proneural transcription factor neurogenin 2 (NGN2; encoded by the *Ngn2*/*Neurog2* gene) has been shown to promote efficient differentiation of mouse ESCs into glutamatergic neurons [44]. In that study, *Ngn2* was expressed from a constitutive promoter following the 4-hydroxytamoxifen induced Cre recombination [44]. Yet, in mice, Ngn2 upregulation in NPCs is transient, followed by a decrease in its expression at later stages of differentiation [45]. To better recapitulate the endogenous regulation, we knocked in a mouse *Ngn2* transgene into the A2Lox ESC line [46] under the doxycycline (Dox) inducible promoter *TRE* (Fig. 1A).

**Figure 1.**
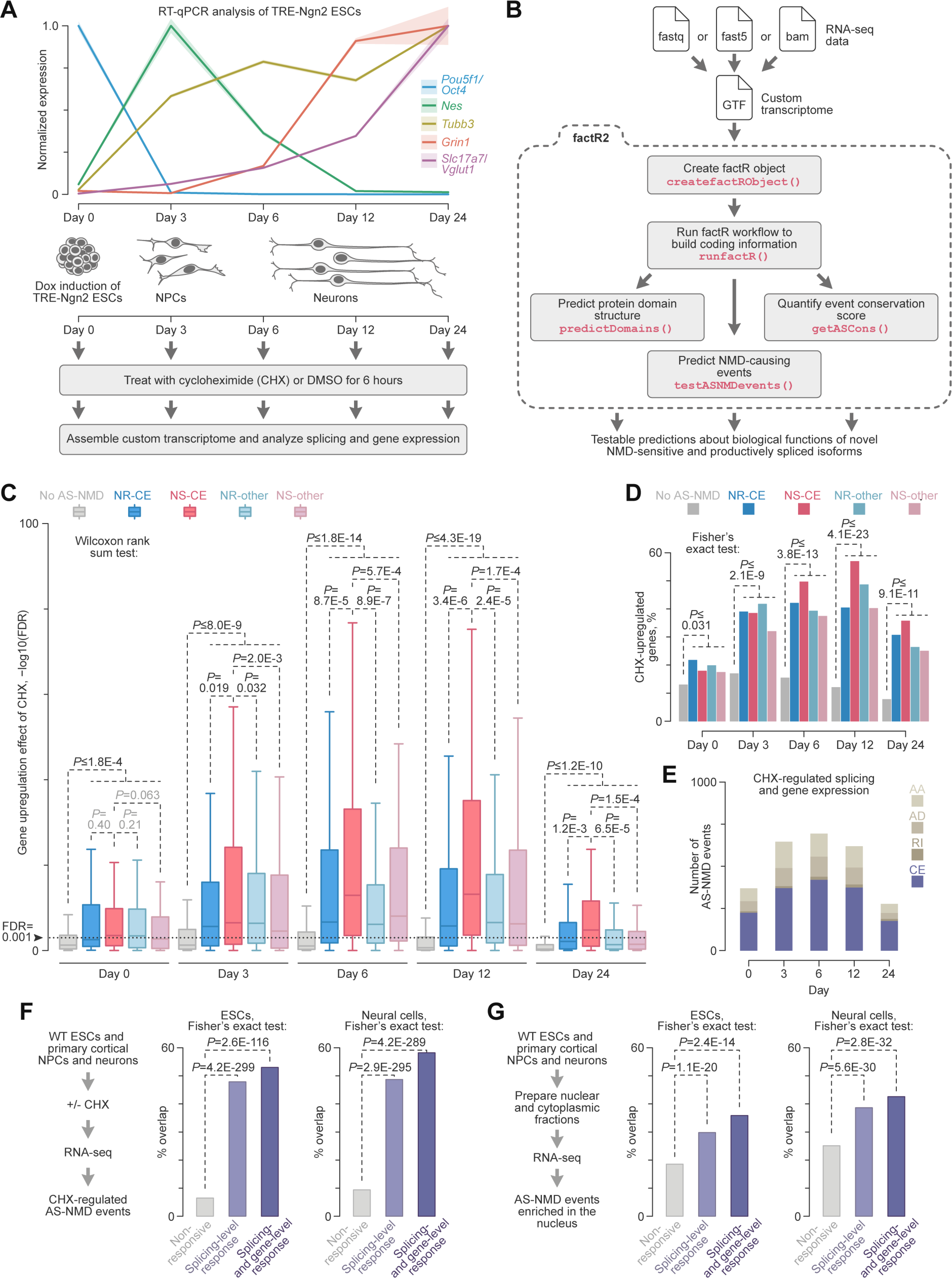
Systematic analysis of the AS-NMD program in developing neurons. (A) Doxycycline (Dox) inducible differentiation of mouse ESCs into neurons. The transition from the ESC to the NPC and neuronal stages is depicted in the *middle* of the panel. *Top*, RT-qPCR analysis of differentiation stage-specific markers in Dox-induced TRE-Ngn2 samples. Note specific expression of the ESC marker *Pou5f1*/*Oct4* on day 0, transient upregulation of the NPC marker *Nes*/*Nestin* on day 3, expression of the pan-neuronal marker *Tubb3* in day 6-24 samples, and upregulation of mature glutamatergic neuron markers *Grin1* and *Slc17a7*/*Vglut1* by days 12 and 24. Data are averaged from 3 experiments with SD shown by the shaded areas. *Bottom*, Dox-induced TRE-Ngn2 cells were treated for 6 hours with DMSO or CHX on days 0, 3, 6, 12, and 24 and analyzed by RNA-seq. (B) Our R package, factR2, streamlines the discovery of functional AS-NMD events and provides a suite of additional tools for functional annotation of custom transcriptomes. (C) Genes containing Whippet-shortlisted AS-NMD events show a stronger upregulation by CHX compared to genes lacking AS-NMD events. In neural cells (days 3-24), this trend is particularly prominent for genes containing NMD-stimulating cassette exons (NS-CEs). The -log10-transformed FDR values for CHX-upregulated genes are presented as a box plot with significance tested by two-tailed Wilcoxon rank sum test. Dotted line, FDR=0.001 value used as a gene regulation significance cutoff throughout this study. (D) Genes containing Whippet-shortlisted factR2 AS-NMD events are more likely to be significantly upregulated (FDR<0.001) by CHX than genes lacking such events. (E) Distribution of AS-NMD events responding to CHX at both the splicing and gene expression levels at different stages of neuronal differentiation. (F) “Natural” samples including wild-type (WT) mouse ESCs (IB10) and NPCs and neurons from mouse embryonic cortex were treated with CHX or DMSO and analyzed by RNA-seq as in (A). CHX-responsive AS-NMD events detected in the natural samples were intersected with three groups of the TRE-Ngn2 factR2 events: not regulated by CHX, regulated by CHX at the splicing level, and regulated by CHX at both the splicing and gene expression levels. The ESC and neural samples were analyzed separately. Note that the natural events are particularly enriched in the TRE-Ngn2 group passing both the splicing and gene regulation cutoffs. (G) Nuclear and cytoplasmic fractions of untreated natural samples introduced in (F) were analyzed by RNA-seq and compared by Whippet. We then calculated the incidence of putative AS-NMD events enriched in the nucleus compared to the cytoplasm among the three TRE-Ngn2 groups mentioned in (F). Again, the largest overlap was detected for TRE-Ngn2 events passing both the splicing and gene regulation cutoffs.

The resultant TRE-Ngn2 line maintained its ESC properties in the 2i+LIF medium [47] without Dox, producing dome-shaped colonies positive for the POU5F1/OCT4 marker (Fig. S1A). When treated with Dox (see Materials and Methods for details), most cells lost OCT4, acquired spindle-shaped morphology typical for NPCs, and increased the expression of the NPC marker vimentin (VIM) by differentiation day 3 (Fig. S1B). To mimic transient upregulation of *Ngn2* in neurogenesis, we progressively reduced Dox concentration beginning from day 2 (see Materials and Methods). By differentiation day 6, most cells stained positive for the early neuronal marker tubulin βIII (TUBB3/TUJ1) and developed characteristic neuronal morphology with round somas and readily detectable neurites (Fig. S1C).

The differentiating cultures were viable for at least 24 days and showed an increased expression of the late neuronal marker MAP2 by days 12-24 (Fig. S1C). Day-24 cells also showed evidence of neuronal maturation including ankyrin G (ANK3/ANKG) staining of the axon initial segment and punctate distribution of the postsynaptic density protein-95 (DLG4/PSD-95) in dendrites (Fig. S1D-E). Reverse transcriptase-quantitative PCR (RT-qPCR) assays confirmed stage-specific expression of corresponding mRNA markers (Fig. 1A). Consistent with the transient Dox treatment used in our protocol, *Ngn2* expression peaked at day 3 and declined at the later stages of differentiation (Fig. S1F).

To understand the role of AS-NMD in neurodevelopment, we treated differentiating TRE-Ngn2 cultures for 6 hours prior to RNA extraction with cycloheximide (CHX), to repress NMD, or DMSO, as a control (Fig. 1A). The samples collected on days 0, 3, 6, 12 and 24 were then analyzed by RNA-seq. Principal component analysis (PCA) of gene expression in this experiment showed a tight clustering of biological replicates and a clear separation of samples according to the differentiation stage and the treatment (Fig. S2A). Further inspection of the DMSO samples confirmed stage-specific expression of pluripotency markers on day 0, NPC markers on day 3, and progressive upregulation of mature neuronal and glutamatergic markers between days 6 and 24 (Fig. S2B). This developmental trajectory was further confirmed by deconvolving cell types in the DMSO samples using MuSiC [48] (Fig. S2C).

We concluded that transient expression of transgenic NGN2 in mouse ESCs provides a suitable system for time-resolved analyses of AS-NMD targets in developing neurons.

### A bioinformatics tool for annotating AS-NMD events in custom transcriptomes

To enable in-depth analysis of AS-NMD targets, we updated our prototype R tool factR (<u>f</u>unctional <u>a</u>nnotation of <u>c</u>ustom <u>tr</u>anscriptomes; [49]) with a suite of relevant functions. The new package, factR2 (Fig. 1B; https://github.com/f-hamidlab/factR2), allocates transcripts identified by RNA-seq to their genes of origin and identifies NMD-sensitive isoforms. It further detects splicing events underlying the NMD sensitivity and provides functions for assessing their regulation potential. factR2 can also deduce domain structure of productively spliced protein-coding sequences and plot RNA and protein isoform structures for data exploration.

As a proof of principle, we searched for NMD-sensitive transcripts among the 90,159 mRNAs present in the GENCODE M26 reference transcriptome using factR2’s predictNMD function. The workflow was kept blind to the GENCODE transcript biotype annotation. This returned 13,042 putative NMD targets containing premature termination codons >50 nt upstream of an exon-exon junction. The set included 7161 out of the 7201 transcripts (i.e. 99.4%) with the “nonsense_mediated_decay” biotype, as well as 5881 additional transcripts that had not been previously flagged as possible NMD targets.

HISAT2-StringTie [50] analysis of our DMSO- and CHX-treated TRE-Ngn2 RNA-seq data (Fig. 1A) predicted 57,535 novel mRNA isoforms absent from the GENCODE annotation. Of these, 30,228 transcripts were classified by factR2 as NMD-sensitive. The analysis of the combined NMD-sensitive transcriptome using factR2’s testASNMDevents function revealed 19,198 putative AS-NMD events. These events were classified (Fig. S3) as cassette exons (CEs), alternative donors (ADs), alternative acceptors (AAs), and retained introns (RIs) that either stimulate or repress NMD when included into mature mRNA (NS and NR events, respectively).

These data demonstrate the utility of factR2 and suggest that the extent of AS-NMD regulation in developing neurons is substantially greater than currently thought.

### factR2 identifies AS-NMD events with a strong regulation potential

To shortlist high-quality AS-NMD targets, we first examined changes in their splicing patterns and gene expression in response to CHX. We used Whippet [51] for the splicing-level analysis and selected CEs, ADs, AAs and RIs consistently changing their percent spliced in (PSI) values in response to CHX at least at one differentiation time point (NS events, ΔPSI>0.1; NR events, ΔPSI<-0.1; probability>0.9). We additionally selected NS events with a PSI average of >0.9 for DMSO- and CHX-treated samples, and NR events with a PSI average of <0.1. This was done because these groups have a limited capacity to change splicing in response to CHX but may still be regulated at the gene expression level.

Genes with Whippet-shortlisted events were more frequently upregulated in response to CHX compared to genes lacking AS-NMD events (Fig. 1C). In many cases, the false discovery rate (FDR) of this effect, calculated by the DESeq2 Wald test [52], was <0.001. Cassette exons stimulating NMD upon their inclusion (NS-CEs) fared particularly well in this analysis in neural cells (days 3-24; Fig. 1C). Notably, the fractions of Whippet-shortlisted entries passing the FDR<0.001 gene-level regulation cutoff were significantly larger compared to genes lacking factR2-predicted AS-NMD events (Fig. 1D). Overall, we identified 1,432 non-redundant AS-NMD events responding to CHX at both splicing and gene levels (Table S1). The number of such events peaked at the intermediate stages of differentiation (days 3-12), with CEs dominating the distributions throughout the entire time course (Fig. 1E). Notably, 49.2% of the CHX-responsive events were not previously annotated in the GENCODE transcriptome.

This analysis suggests that the AS-NMD pathway controls hundreds of genes expressed at different stages of neuronal differentiation.

### Many AS-NMD events identified in TRE-Ngn2 cells are regulated in natural samples

To validate our approach, we performed RNA-seq analyses of wild-type (WT) mouse ESCs (IB10 line) and primary cortical NPCs and neurons treated with CHX or DMSO (Fig. 1F and Fig. S4A-C). Splicing events responding to CHX in these “natural” samples (with the Whippet cutoffs described above) were significantly overrepresented among the CHX-responsive TRE-Ngn2 factR2 events compared to the non-responsive TRE-Ngn2 factR2 control (Fig. 1F).

Since NMD occurs in the cytoplasm [12–14], transcripts including NMD-stimulating and excluding NMD-repressing events should be enriched in the nucleus, providing a CHX-independent validation approach. With this in mind, we sequenced nuclear and cytoplasmic transcriptomes from untreated WT ESCs and cortical NPCs and neurons, and compared the two compartments using Whippet (Fig. 1G). Nucleus-enriched NR and nucleus-depleted NS events were overrepresented in the CHX-responsive TRE-Ngn2 factR2 sets (Fig. 1G). Notably, the sequence context of events responding to CHX in both the TRE-Ngn2 and natural cells was more conserved across vertebrates compared to a non-responsive control (Fig. S4D-E).

These data argue that many AS-NMD events identified by factR2 are regulated in vivo. Since the targets responding to CHX at both the splicing and gene expression levels showed the largest overlap with the natural regulation (Fig. 1F-G) and the highest median conservation scores (Fig. S4D-E), we prioritized this set in our subsequent analyses.

### AS-NMD is widespread among genes downregulated in developing neurons

So far, we focused on the impact of AS-NMD on individual stage-specific transcriptomes. To explore potential effects of this pathway on gene expression dynamics in development, we analyzed the correlation between splicing patterns protecting mRNA from NMD (i.e. inclusion of NR events and skipping of NS events) and gene expression levels. This was done by applying factR2’s function testGeneCorr to genes differentially expressed (DESeq2 likelihood-ratio test; FDR<0.001) in the control-treated longitudinal TRE-Ngn2 data.

Compared to the non-responsive control, CHX-responsive AS-NMD events were enriched for positively correlated entries (Fig. 2A). The enrichment was predominantly due to CEs rather than other types of AS-NMD events (Fig. 2A-B). The largest fraction of significantly correlated events was detected for NS-CEs, with the magnitude of this effect increasing for tighter *P*-value cutoffs (Fig. 2A-B). Hereafter, we will use the term “facilitating” to refer to CHX-responsive events showing significant positive correlation (*P*<0.05) between NMD-protective splicing patterns and gene expression.

**Figure 2.**
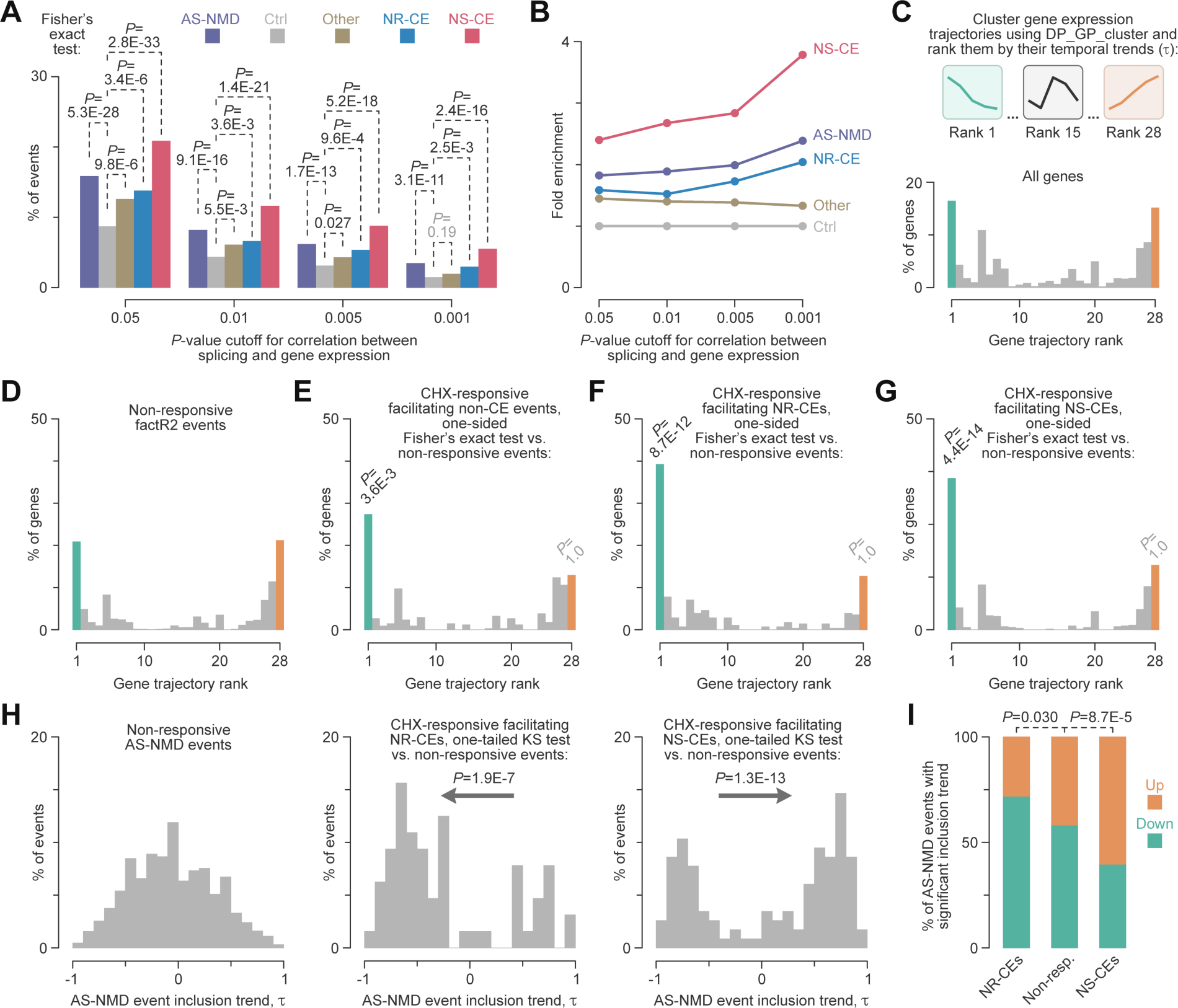
Neurodevelopmentally downregulated genes are often controlled by AS-NMD. **(A-B)** factR2 events responding to CHX at both the splicing and gene levels show a stronger correlation with developmental changes in gene expression compared to non-responsive events. (A) Percentages of events correlating with gene expression dynamics in differentiating TRE-Ngn2 cells for different *P*-value cutoffs. Ctrl are factR2 events not responding to CHX. AS-NMD, Other, NR-CE, and NS-CE are, respectively, all AS-NMD events, AD, AA and RI AS-NMD events, NMD-repressing cassette exons, and NMD-stimulating cassette exons responsive to CHX. (B) Fold enrichments of the correlated events in the shortlisted categories in (A) plotted as a function of the correlation *P*-value cutoff. NS-CEs show the strongest enrichment for correlated entries, with the fold enrichment increasing for tighter cutoffs. **(C)** *Top*, gene expression trajectory analysis in differentiating TRE-Ngn2 cells. *Bottom*, distribution of ranked trajectories for all regulated genes (DESeq2’s LRT test, FDR<0.001). **(D-G)** Ranked trajectories for regulated genes containing (D) only non-responsive factR2 events; (E) CHX-responsive facilitating AD, AA and RI events; (F) CHX-responsive facilitating NR-CEs; (G) CHX-responsive facilitating NS-CEs. The rank-1 (monotonic downregulation) and rank-28 (monotonic upregulation) trajectories were compared to the non-responsive AS-NMD control by one-sided Fisher’s exact test. Note that monotonic downregulation is particularly prevalent in (F-G). (H) Neurodevelopmental changes in AS-NMD event inclusion status calculated as Kendall’s τ for PSI trend. Note significant overrepresentation of the decreasing trend among CHX-responsive facilitating NR-CEs (middle) and the increasing trend among CHX-responsive facilitating NS-CEs (right), compared to non-responsive AS-NMD events (left). The *P*-values were calculated by one-sided Kolmogorov-Smirnov (KS) test. (I) Preferential decrease in NR-CE inclusion and increase in NS-CE inclusion in developing neurons is also detectable by Fisher’s exact test analysis of events with statistically significant Kendall’s trends (*P*<0.05).

Since the above results pointed to a possible role of AS-NMD in gene expression dynamics, we wondered if such a mechanism might preferentially feed into specific types of temporal regulation. We clustered differentially expressed genes (DESeq2 likelihood-ratio test; FDR<0.001) according to their TRE-Ngn2 time-course trajectories using DP_GP_cluster [53]. We then ranked the 28 trajectories identified by this analysis based on the τ statistic of the Kendall’s trend analysis, from monotonic downregulation (rank 1) to monotonic upregulation (rank 28) (Fig. 2C, Fig. S5, and Table S2).

Strikingly, the facilitating AS-NMD events were often associated with the rank-1 downregulation trajectory (Fig. 2D-G and Table S3). The enrichment was especially robust for genes with facilitating NS-CEs (one-sided Fisher’s exact test, *P*=4.4E-14; Fig. 2G). Similar trends were detected when we analyzed the trajectories of individual genes without clustering (Fig. S6A-C). Consistent with the opposite effects of NR-CE and NS-CE inclusion on mRNA stability, Kendall’s analysis of the PSI dynamics revealed a skew towards negative τ values in NR-CEs and positive τ values in NS-CEs (Fig. 2H). Events with significant positive PSI trends (*P*<0.05) were significantly overrepresented in the NS-CE group (Fisher’s exact test, P=8.7E-5; Fig. 2I). Moreover, the intronic sequence context of NS-CEs encoded in monotonically downregulated genes showed a stronger interspecies conservation compared to other groups of CEs (Fig. S6D-E).

Overall, our pipeline identified 87 facilitating NS-CEs associated with monotonic gene downregulation (Kendall’s τ<-0.75, *P*<0.05; Table S3). Metascape [54] analysis of this set revealed significant enrichment for RNA metabolism and localization as well as cell cycle regulation functions (Fig. S7A and the first tab in Table S4). Other examples of the overrepresented functional categories included RHO GTPase effectors and actin cytoskeleton organization (Fig. S7A).

To validate our bioinformatics predictions, we examined four monotonically downregulated genes (τ<-0.75, *P*<0.05; Table S3) with evolutionarily conserved NS-CEs (Fig. S8). These included three examples encoding RNA metabolism factors, *Fbl*, *Srsf9*, and *Xpo1* (the first tab in Table S4), and the *Ctnnal1* gene predicted to contribute to actin filament binding and RHO signaling (https://www.ncbi.nlm.nih.gov/gene/8727). All four NS-CEs were readily detectable in CHX-treated samples showing progressively stronger inclusion at later stages of differentiation.

Thus, AS-NMD in general and NMD-stimulating cassette exons in particular may play a role in downregulation of functionally related groups of genes in developing neurons.

### Comprehensive identification of AS-NMD events regulated by PTBP1

Previous studies by our group and others identified several AS-NMD events controlled by PTBP1, an RNA-binding protein expressed at high levels in ESCs and NPCs and downregulated in neurons [28–31, 34]. We therefore used PTBP1-regulated events to benchmark the performance of the factR2-based workflow. *Ptbp1* was monotonically downregulated in differentiating TRE-Ngn2 cells, and the short list of facilitating AS-NMD events included previously characterized PTBP1 targets (e.g., NR-CEs maintaining open reading frames in *Ptbp2*, *Gabbr1* and *Dlg4*; Fig. S9A and Table S3).

To identify PTBP1-dependent AS-NMD targets systematically, we examined factR2 events correlating with *Ptbp1* expression in DMSO-treated TRE-Ngn2 cells and changing their splicing in ESCs or NPCs in response to PTBP1 knockdown alone or together with its functionally similar paralog, PTBP2 [55] (Whippet knockdown response |ΔPSI|>0.1; probability>0.9 cutoffs; see Materials and Methods for further details; Fig. 3A).

**Figure 3.**
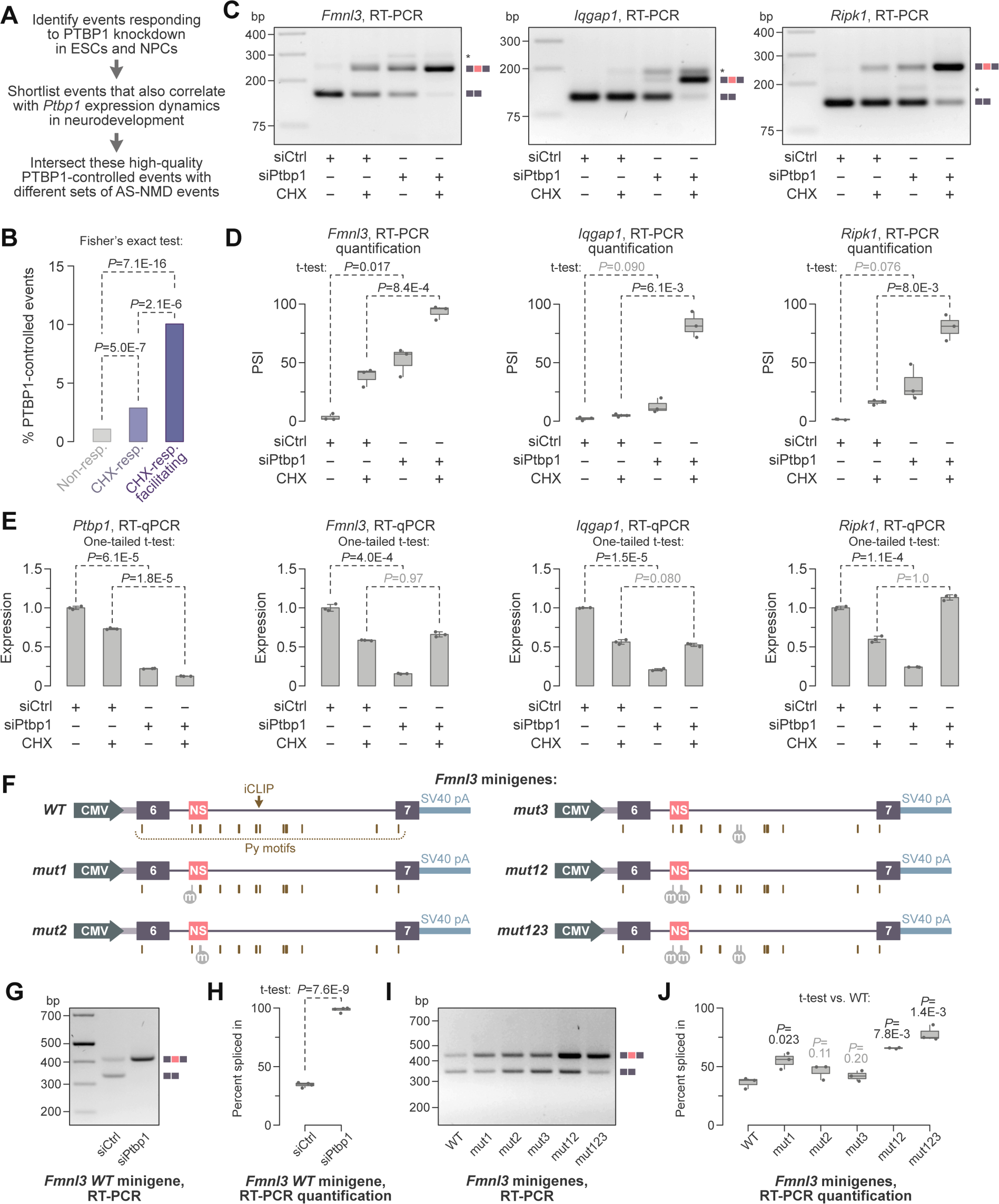
Role of PTBP1 in AS-NMD regulation in developing neurons. **(A)** The workflow used to shortlist PTBP1-controlled events. **(B)** AS-NMD events with strong regulatory potential are enriched for PTBP1-controlled entries. **(C-E)** Uninduced TRE-Ngn2 ESCs were treated for 48 hours with a PTBP1-specific (siPtbp1) or a non-targeting control siRNA (siCtrl) and post-treated with CHX (+CHX) or DMSO (-CHX) for an additional 6 hours. (C) The samples were then analyzed by RT-PCR with primers flanking the *Fmnl3*, *Iqgap1* and *Ripk1* NS-CEs and the amplified products were separated by agarose gel electrophoresis. Note that PTBP1 represses NS-CE inclusion and CHX stabilizes NS-CE-containing splice forms. (D) The NS-CE percent spliced in (PSI) values in (C) were quantified from 3 experiments and shown as box plots. (E) RT-qPCR analyses with primers against constitutively spliced parts of *Fmnl3*, *Iqgap1* and *Ripk1* show that siPtbp1 dampens the expression of these genes in the DMSO but not the CHX samples. We also used *Ptbp1*-specific primers to estimate its knockdown efficiency. The data are averaged from three experiments ±SD. **(F)** *Fmnl3* minigenes used in this study. Pyrimidine-rich (Py) motifs predicted to recruit PTBP1 to the NS-CE or to the intronic positions overlapping with a PTBP1 iCLIP cluster [55] have been mutated (gray circles labeled by “m”) in the *mut1*, *mut2*, *mut3*, *mut12*, and *mut123* minigenes. **(G)** TRE-Ngn2 ESCs pretreated with siPTBP1 or siCtrl were transfected with the wild-type (*WT*) *Fmnl3* minigene and analyzed by RT-PCR with a minigene-specific primer pair. Note that the minigene recapitulates the repressive effect of PTBP1 on the *Fmnl3* NS-CE. **(H)** Quantification of the NS-CE PSI values in (G). **(I)** Untreated TRE-Ngn2 ESCs were transfected with the *Fmnl3* minigenes introduced in (F) and analyzed by RT-PCR. **(J)** Quantification of the PSI values in (I) showing that the mutation of the upstream exonic Py (*mut1*) promotes the NS-CE inclusion, especially when combined with the genetic inactivation of the downstream exonic Py (*mut2*) and the iCLIP Py positions (*mut3*). Data in (H and J) were obtained from 3 experiments, presented as box plots, and compared by two-tailed t-test assuming unequal variances.

Notably, the PTBP1-dependent behavior was significantly overrepresented among the high-quality TRE-Ngn2 factR2 events (Fig. 3B). Of the 46 facilitating AS-NMD events regulated by PTBP1, 29 were NS-CEs (Table S3). Similar to the trend observed for all facilitating NS-CEs (Fig. S6B-C), the PTBP1-regulated NS-CEs were strongly enriched in monotonically downregulated (τ<-0.75, P<0.05) but not upregulated (τ>0.75, P<0.05) genes (Fig. S9B-C).

Metascape analysis showed that monotonically downregulated genes with PTBP1-controlled NS-CEs were enriched for the RHO GTPase effector and actin cytoskeleton organization functions (Fig. S7B and the second tab in Table S4). Another category detected by this analysis related to the programmed cell death (apoptosis modulation by HSP70; Fig. S7B). Interestingly, the overrepresentation of the RHO GTPase effector and actin cytoskeleton organization terms among all downregulated genes with NS-CEs (Fig. S7A) was largely due to the PTBP1-regulated targets. Indeed, these categories were missing in the Metascape output when we analyzed monotonically downregulated genes with PTBP1-independent NS-CEs (Fig. S7C and the third tab in Table S4).

These data suggest that PTBP1 controls a wider range of NS-CEs than previously thought, and that many genes containing these events belong to specific functional categories.

### PTBP1-mediated repression of NS-CEs promotes the expression of their genes in ESCs

We selected three neurodevelopmentally downregulated genes with PTBP1-repressed NS-CEs for further analyses: *Fmnl3* and *Iqgap1* encoding RHO GTPase effectors involved in actin cytoskeleton organization and *Ripk1*, a regulator of programmed cell death (the second tab in Table S4). The three genes were strongly downregulated in developing neurons in a CHX-rescuable manner (Fig. S9D-F). Their NS-CEs were evolutionarily conserved and neighbored or overlapped with PTBP1 interaction motifs (YUCUYY and YYUCUY) and PTBP1 binding sites identified by iCLIP [55] (Fig. S10). PTBP1 was proposed to control *Ripk1* and *Iqgap1* NS-CE splicing [27, 56–58], but the role of these exons in the regulation of their host genes during neuronal differentiation was not addressed experimentally. To the best of our knowledge, the Fmnl3 NS-CEs has not been previously identified.

To validate the role of PTBP1 in the expression of these targets, we transfected TRE-Ngn2 ESCs with Ptbp1-specific or control siRNAs (siPtbp1 or siCtrl, respectively) and post-treated the cells with either CHX or DMSO. Subsequent RT-PCR analyses showed that siPTBP1 stimulated the inclusion of all three NS-CEs, which was further enhanced by CHX (Fig. 3C-D). PTBP1 knockdown dampened the total mRNA abundance for the three genes in the DMSO-but not CHX-treated samples (Fig. 3E). Importantly, the downregulation of *Fmnl3*, *Iqgap1* and *Ripk1* by siPtbp1 was also significantly rescued by an siRNA against the key NMD factor UPF1 (Fig. S11).

To test if PTBP1 might regulate NS-CE splicing directly, we cloned the *Fmnl3* NS-CE in its natural sequence context into a mammalian expression vector (Fig. 3F). Transcripts produced from this minigene lacked an in-frame translation initiation codon and were not predicted to undergo NMD. The wild-type minigene (*WT*) recapitulated the endogenous splicing regulation in TRE-Ngn2 ESCs (Fig. 3G-H). There are two putative PTBP1 binding sites inside the *Fmnl3* NS-CE and two such positions in the downstream intron that overlap with a PTBP1 iCLIP cluster (Fig. S10A). Mutation of the upstream exonic site (*mut1*; Fig. 3F) increased the NS-CE inclusion in the presence of PTBP1 (Fig. 3I-J). Mutations of the downstream exonic site (*mut2*) or the two iCLIP positions (*mut3*) did not alter the minigene splicing pattern individually, but enhanced the NS-CE stimulation effect of *mut1* in the *mut12* and *mut123* minigenes (Fig. 3F,I-J).

We concluded that PTBP1 promotes the expression of several non-neuronal genes in ESCs by repressing NS-CEs.

### Ptbp1 downregulation in developing neurons may act as a switch for NS-CE inclusion

The TRE-Ngn2 RNA-seq data showed a substantial decrease in *Ptbp1* expression between differentiation days 3 (NPCs) and 6 (young neurons), which coincided with the expression trajectories of *Fmnl3*, *Iqgap1* and *Ripk1* (Fig. S9A). To find out whether the *Ptbp1* dynamics could promote the NS-CE inclusion in these targets, we analyzed the TRE-Ngn2 differentiation time course collecting DMSO-or CHX-treated samples daily between days 0 and 6 (Fig. 4).

**Figure 4.**
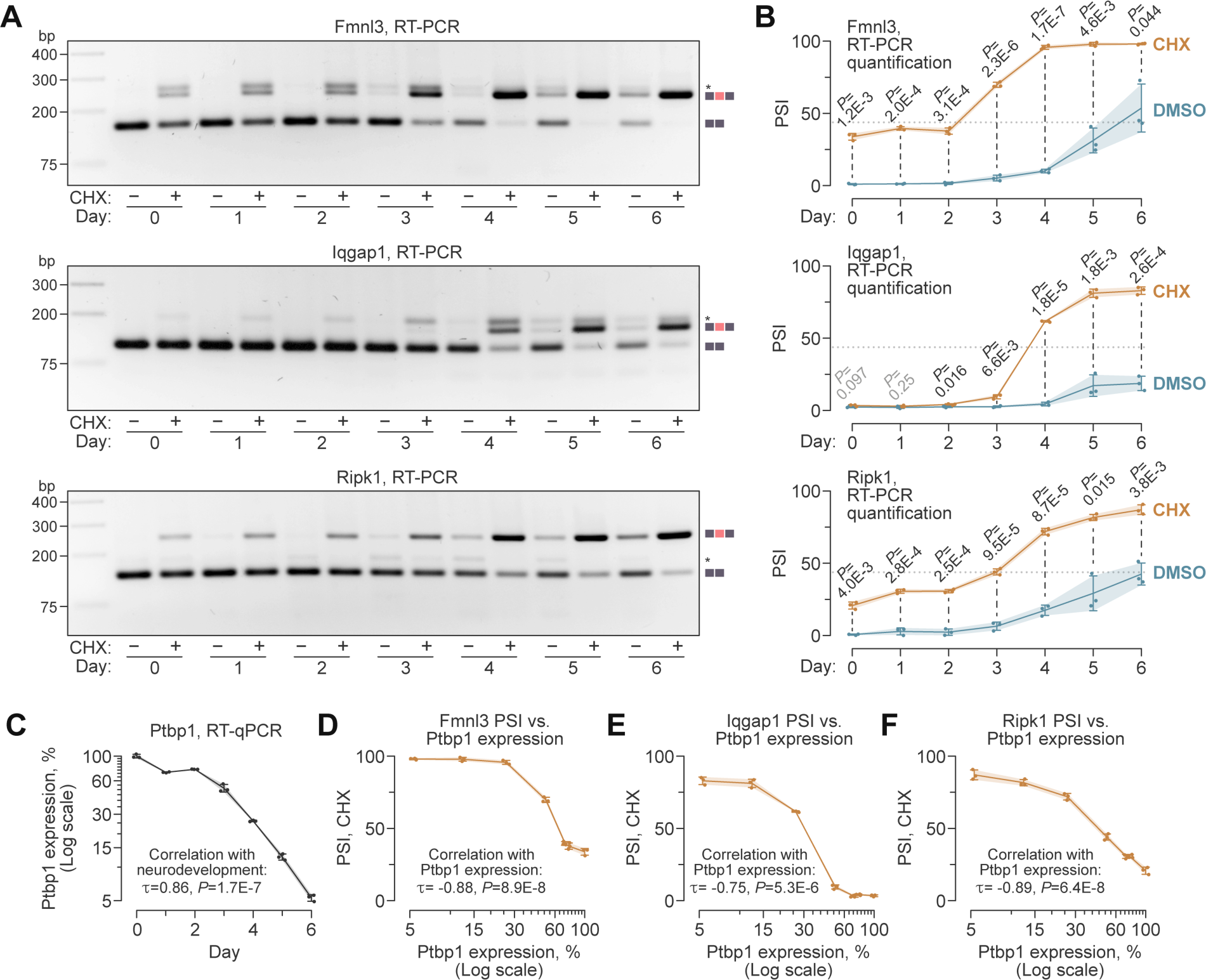
PTBP1-repressed NS-CEs are regulated in a switch-like manner during neuronal differentiation. **(A)** RT-PCR analyses of Dox-induced TRE-Ngn2 ESCs treated with DMSO or CHX on days 0-6 show that the inclusion of NS-CEs into the Fmnl3, Iqgap1 and Ripk1 mRNAs increases during neuronal differentiation. **(B)** NS-CE PSI values for the DMSO and CHX samples in (A) averaged from three experiments ±SD and compared by two-tailed t-test assuming unequal variances. Note that the CHX curves are S-shaped, consistent with a switch-like regulation. Dotted lines, PSI median of the CHX-treated data for the three genes used to estimate the onset of the rapid increase phase. **(C)** RT-qPCR analysis Ptbp1 expression dynamics during the day 0-6 differentiation period. Note that the abundance of the Ptbp1 mRNA decreases monotonically beginning from day 2, consistent with the onset of the rapid increase phase in the CHX curves in (B). **(D-F)** NS-CE inclusion in (D) Fmnl3, (E) Iqgap1, and (F) Ripk1 shows a strong negative correlation with Ptbp1 expression, as expected. Correlations in (C-F) were analyzed using the Kendall’s method.

RT-PCR analysis of DMSO-treated samples (Fig. 4A) did not detect the NS-CE-containing isoforms of Fmnl3, Iqgap1 and Ripk1 between days 0 and 2. Small amounts of these isoforms appeared on days 3-4 and accumulated at the subsequent stages of differentiation. Treating the cells with CHX tended to increase the relative abundance of the NS-CE-containing isoforms with a robust difference between the CHX and the DMSO PSI in newly developed neurons (*P*<1E-4; Fig. 4B).

Since CHX protects NS-CE-containing isoforms from NMD, we used the CHX-treated data to follow developmental changes in pre-mRNA splicing patterns (Fig. 4B). In all three cases, the time-course curves were S-shaped with little or no changes in the NS-CE inclusion between days 0-2, a rapid increase between days 2-4, and a plateau between days 4-6. Using the median PSI value for the three genes as a threshold (43.8%; dotted line in Fig. 4B) placed the onset of the rapid increase phase at day 2 for *Fmnl3* and day 3 for *Iqgap1* and *Ripk1*. RT-qPCR analysis of *Ptbp1* expression showed that it was relatively stable between days 0-2, and progressively declined beginning from day 2 (Fig. 4C). Notably, NS-CE-inclusion values showed a strong negative correlation with *Ptbp1* for all three targets (Fig. 4D-F).

Taken together, these results suggest that reduced *Ptbp1* expression may facilitate a switch towards the NS-CE inclusion during the transition from NPCs to neurons.

### NS-CEs work in coordination with NMD-independent gene downregulation mechanisms

Increased inclusion of NS-CEs may contribute to gene downregulation in developing neurons in four different ways. (1) It may act as the sole downregulation mechanism. Alternatively, it may work alongside AS-NMD-independent repression mechanisms by (2) preempting, (3) coinciding with, or (4) following them in development. We modeled these four scenarios assuming possible cooperation with transcriptional repression (Fig. S12 and Materials and Methods). The key predictions from this simple model were that AS-NMD should consistently diminish residual gene expression later in development in all four scenarios and accelerate the onset of downregulation in scenarios (1) and (2), but not (3) and (4).

To find out which of the above mechanisms could be used in developing neurons, we targeted NS-CE sequences in *Fmnl3*, *Iqgap1* and *Ripk1* using appropriate CRISPR gRNAs and Cas9 (Fig. S13). For each gene, we selected two TRE-Ngn2 ESC clones with biallelic NS-CE disruption (Fig. S13). RT-PCR analyses of undifferentiated (day-0 ESCs) and Dox-differentiated (day-6 neurons) samples treated with DMSO or CHX showed that the mutant clones lost or had a severely impaired ability to splice in NMD-stimulating exons (Fig. S14).

We then analyzed the expression dynamics of the three genes in the wild-type (WT) and mutant cells undergoing neuronal differentiation (Fig. 5A-C). As expected, the three WT genes were downregulated in day-6 neurons compared to day-0 ESCs. *Iqgap1* and *Fmnl3* additionally showed transient upregulation peaks on days 1-2, which could not be detected in our day 0, 3, 6, 12 and 24 RNA-seq time series. The mutants generally followed the WT trends (Fig. 5A-C), with two key differences revealed by normalizing the data by the wild-type values (Fig. 5D-F).

**Figure 5.**
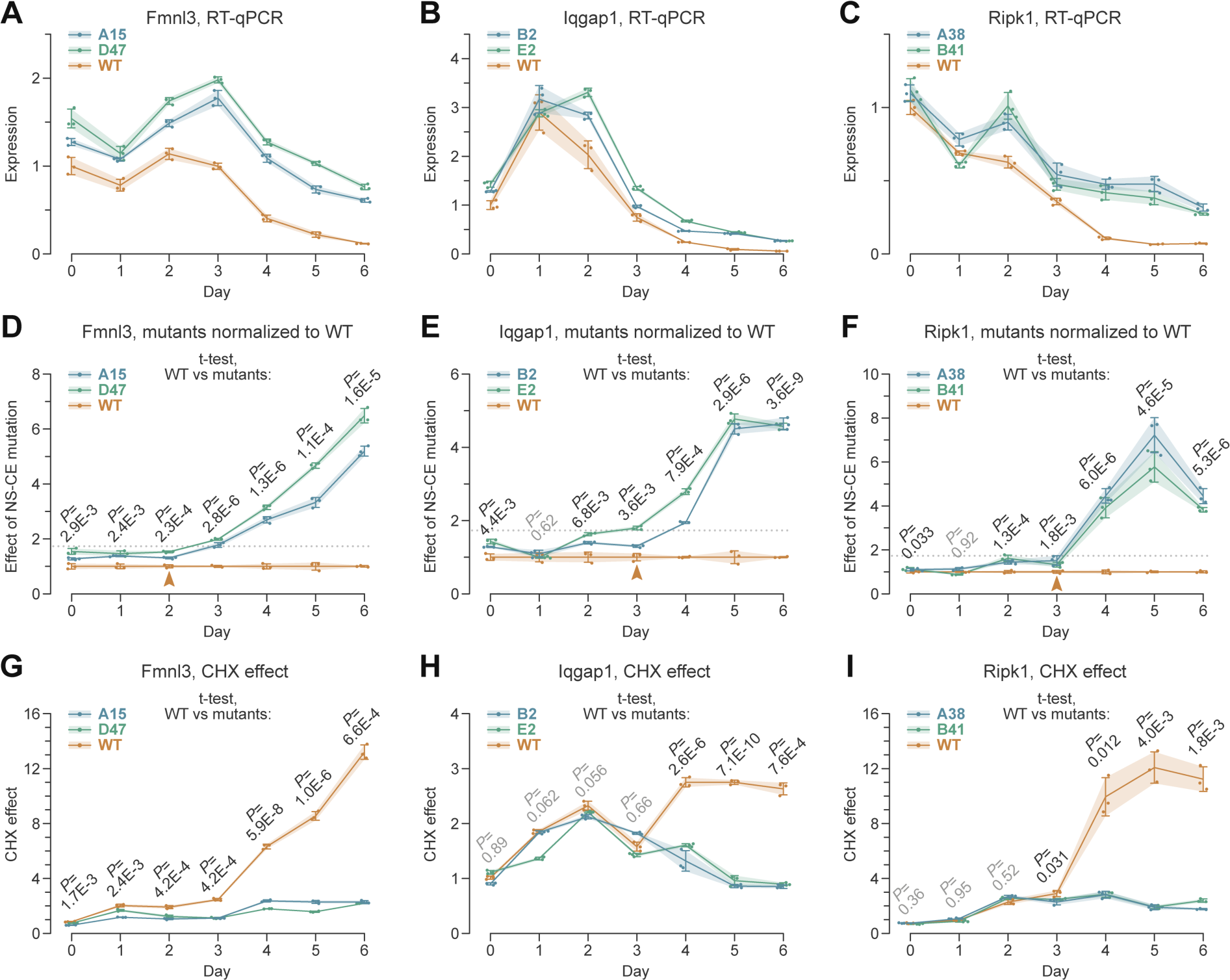
NS-CEs facilitate neurodevelopmental downregulation of non-neuronal genes. **(A-C)** TRE-Ngn2 ESCs with wild-type (WT) or biallelically mutated NS-CEs (*Fmnl3*, clones A15 and D47; *Iqgap1*, clones B2 and E2; and *Ripk1*, clones A38 and B41) were differentiated into neurons and the gene expression dynamics was analyzed by RT-qPCR. Note that the mutant (A) *Fmnl3*, (B) *Iqgap1* and (C) *Ripk1* have a higher residual expression in neurons compared to their WT counterparts. Mutant *Fmnl3* is also expressed somewhat higher than the WT at earlier stages of differentiation. **(D-F)** Normalizing the data in (A-C) to the WT values confirms that (D) *Fmnl3*, (E) *Iqgap1* and (F) *Ripk1* with mutated NS-CEs have a substantially reduced ability to undergo downregulation at later stages of development. We defined WT-mutant dichotomy points as differentiation stages preceding the increase of normalized expression values averaged for the two mutants above the median of all normalized mutant values calculated for the three genes (1.73; dotted line). This places the dichotomy points at day 2 for Fmnl3 and day 3 for Iqgap1 and Ripk1 (arrowheads). **(G-I)** Normalizing CHX-treated time-course to the corresponding DMSO-treated controls for (G) *Fmnl3*, (H) *Iqgap1* and (I) *Ripk1* confirms that the AS-NMD regulation is lost or markedly diminished in the NS-CE mutants, and that days 2-3 correspond to the onset of facilitating AS-NMD in the WT.

*First*, NS-CE inactivation led to progressive upregulation of gene expression at later developmental stages (Fig. 5D-F). For all 3 genes, the mutant trajectories diverged from their WT counterparts after an initial lag period. By using the median of the normalized mutant values for all three genes as a cutoff (1.73; dotted lines in Fig. 5D-F), we estimated that the WT-mutant dichotomy point was around day 2 for *Fmnl3* and day 3 for Iqgap1 and Ripk1. According to Fig. S12, these points mark the onset of facilitating AS-NMD in the WT, and they indeed coincided with the corresponding parts of the NS-CE inclusion curves in CHX-treated WT samples (Fig. 4B).

*Second*, the temporal relationship between the AS-NMD dichotomy points in Fig. 5D-F and the monotonically downregulated parts of the mutant trajectories in Fig. 5A-C varied depending on the gene. The estimated onset of AS-NMD preceded the monotonic downregulation for the *Fmnl3* (day 2 vs. day 3), while following it for *Iqgap1* and *Ripk1* (day 3 vs. day 2). These behaviors matched models (2) and (4) in Fig. S12, with apparent one-day difference between the onset of NMD-dependent and -independent downregulation mechanisms. Confirming the role of the NS-CEs in the AS-NMD regulation, CHX strongly upregulated the WT but not the mutant genes at the corresponding differentiation stages (Fig. 5G-I).

Thus, in the three examples tested, NS-CEs are essential for full-scale downregulation of non-neuronal genes in developing neurons, working in coordination with NMD-independent repression mechanisms.

### NS-CEs may contribute to gene downregulation in other developmental contexts

The NS-CEs investigated above were identified in developing glutamatergic neurons. To examine the incidence of NS-CEs in a different system, we performed factR2 analysis for developing dentate gyrus granule cells [59], which are known to secrete a combination of neurotransmitters [60]. Since the granule cell dataset lacked the CHX treatment, we used facilitation (i.e. a positive correlation between splicing patterns protecting mRNA from NMD and gene expression) as the sole criterion to shortlist 235 NS-CE candidates (Table S5). The rationale for this approach was that CHX-responsive NS-CEs were enriched for the facilitating behavior in our TRE-Ngn2 analysis (Fig. 2A-B). Consistent with the cell type-specific differences in gene expression, many of the granule cell events were novel compared to their TRE-Ngn2 counterparts. An interesting example of the shared targets was the Bak1 NS-CE previously shown to dampen the expression of this pro-apoptotic factor in developing neurons [32].

Strikingly, nearly 60% of the granule cell genes containing shortlisted NS-CEs were monotonically downregulated during granule development (Kendall’s τ<-0.75, *P*<0.05). This constituted a ∼4.3-fold enrichment (one-sided Fisher’s exact test, *P*=1.8E-57) compared to a control group containing CEs not associated with AS-NMD (Fig. 6A). We did not detect such enrichment for monotonically upregulated genes (Fig. 6B). Notably, intronic sequence context of NS-CEs in monotonically downregulated but not upregulated genes showed a stronger interspecies conservation compared to non-AS-NMD CEs (Fig. 6C-D).

**Figure 6.**
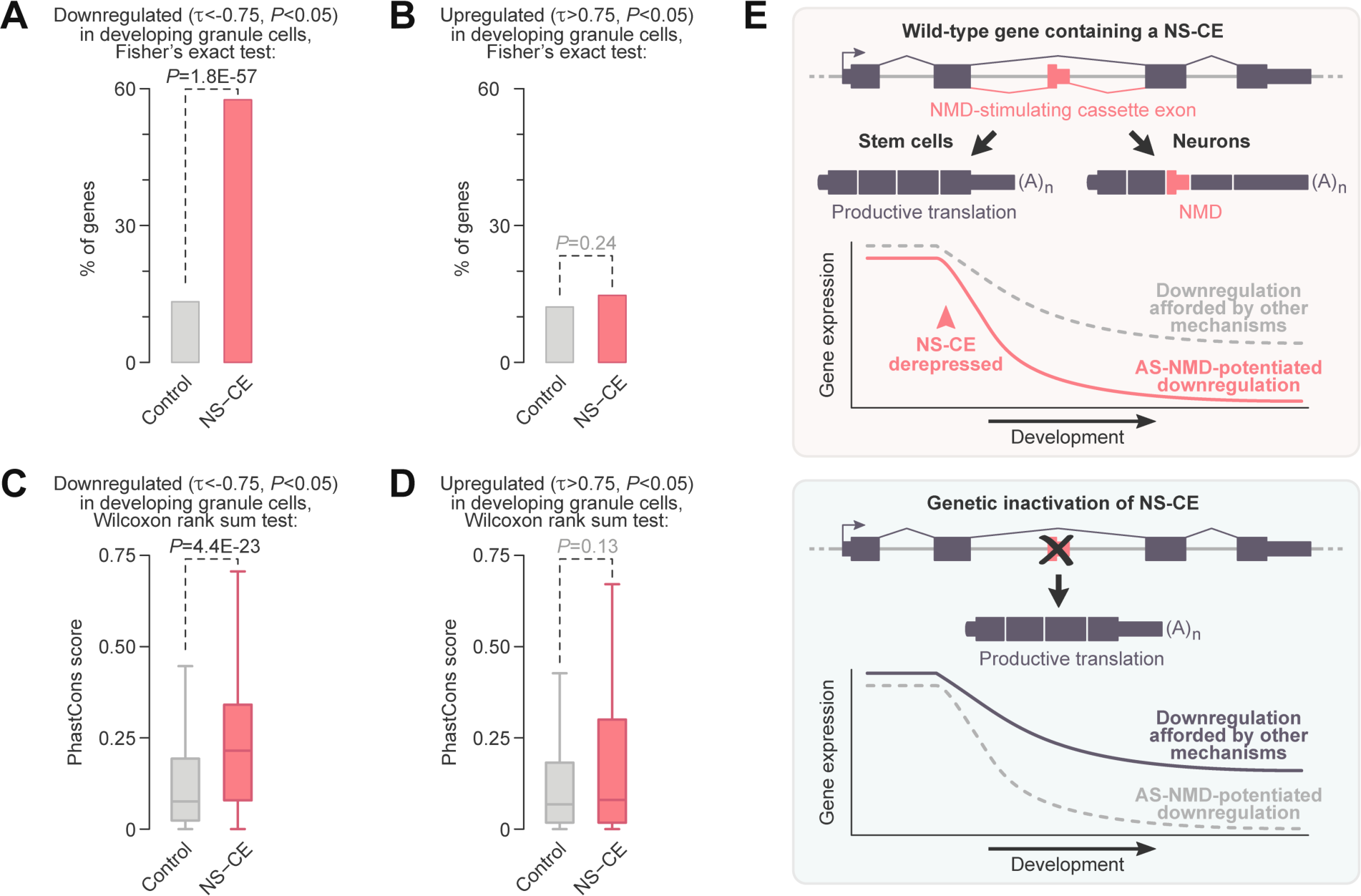
NS-CEs provide a widespread mechanism potentiating gene downregulation in developing neurons. **(A-B)** High-quality NS-CE events identified by factR2 analysis of developing dentate gyrus granule cells [59] are enriched in developmentally downregulated (A) but not upregulated (B) genes compared to CE-containing genes not predicted by factR2 to undergo AS-NMD. **(C-D)** The granule-cell NS-CEs encoded in developmentally downregulated, but not upregulated genes tend to be conserved [94] significantly stronger compared to the corresponding groups of CEs not predicted by factR2 to be involved in AS-NMD. **(E)** Model – see text for details.

These analyses indicate that NS-CEs are widespread and evolutionarily conserved regulatory elements involved in gene downregulation in developing neurons.

## Discussion

Our study provides valuable insights into the gene regulation functions of the AS-NMD pathway. Firstly, it introduces a straightforward workflow for shortlisting biologically relevant AS-NMD targets. Secondly, it reveals the extensive involvement of AS-NMD in downregulation of non-neuronal genes in developing neurons. Finally, it argues that the activity of NMD-stimulating cassette exons (NS-CEs) is often coordinated with other downregulation mechanisms to ensure full-scale repression of their target genes in mature neurons (Fig. 6E).

Our optimized inducible differentiation system provides a useful resource for time-resolved analyses of gene expression dynamics in developing neurons (Fig. 1A, Fig. S1 and Fig. S2). We also believe that users working with custom transcriptomes deduced from either short- or long-read sequencing data will appreciate the ability of factR2 (Fig. 1B) to quickly allocate sample-specific transcripts to their original genes (done when creating the factR2 object), predict open reading frames (buildCDS), identify transcripts containing NMD-promoting features (predictNMD), and pinpoint AS events underlying NMD sensitivity (testASNMDevents). Functions testGeneCorr and getAScons are helpful for assessing the regulatory potential and evolutionary conservation of splicing events, while plotTranscripts provides rich data visualization capabilities. Although not showcased in this study, factR2 can be also used to predict the domain structure of proteins encoded by productively spliced isoforms (predictDomain).

In our evaluation, factR2 outperforms its earlier alternatives [61, 62] in terms of portability, the range of functions, and seamless integration with other R packages used for bioinformatics analyses. The efficiency of factR2 as an AS-NMD discovery tool is supported by its ability to identify the vast majority of the GENCODE-annotated NMD targets (99.4%). Moreover, when applied to a custom transcriptome containing both reference and novel isoforms, factR2 expanded the NMD biotype annotation by ∼6-fold (43,270 vs. 7,201).

factR2 analysis of TRE-Ngn2 neuronal differentiation time series identified hundreds of AS-NMD targets responding to the NMD inhibitor CHX at specific time points (Table S1 and Fig. 1E). Confirming their biological relevance and authenticity, many of these events were also detected in genetically unperturbed ESCs and ex vivo neural cells (Fig. 1F-G). Interestingly, the number of CHX-responsive targets increased transiently in NPCs and developing neurons (days 3-12; Fig. 1E), suggesting that AS-NMD might be particularly important at differentiation stages associated with a large-scale rewiring of the transcriptome. The relative paucity of AS-NMD targets in mature neurons (day 24) is consistent with previously reported downregulation of the NMD factors, UPF1 and MLN51 by the neuronal microRNAs [39, 40].

Our data corroborate earlier findings that a considerable fraction of mammalian genes generate NMD-sensitive isoforms [14, 26, 62–64]. A key advancement of the present study is the identification of events with a strong potential to modulate gene expression during development. By focusing on this high-quality subset, we show that AS-NMD events, and their NS-CE subset in particular, are enriched in genes downregulated during neuronal differentiation (Fig. 2, Fig. S6 and Table S3). Our reanalysis of a published granule cell dataset [59] argues for a broader association between NS-CEs and gene downregulation in developing neurons (Fig. 6A-D and Table S5).

This is an important finding since previously characterized examples of exonic AS-NMD events have shown varied effects on the expression of their host genes, including both up- and downregulation in developing neurons [28–34]. The regulation of *Ptbp2* follows a more complex pattern with initial upregulation in young neurons and subsequent downregulation during neuronal maturation [31]. Whether any of these scenarios are favored at the systems level has been an open question. In contrast, a post-transcriptional circuitry operating in developing granulocytes involves a coordinated increase in intron retention in 86 functionally related genes, dampening their expression in an NMD-dependent manner [22].

We detected 87 NS-CEs in genes monotonically downregulated in differentiating TRE-Ngn2 cells and enriched for specific biological functions (Fig. S7A and Tables S3 and S4). Thus, despite the obvious difference in the type of regulated splicing events, AS-NMD provides a transcriptome-wide mechanism for repressing functionally related genes in both neuronal and granulocyte development. The sequence context of the facilitating NS-CEs identified by our pipeline showed a remarkable degree of evolutionary conservation (Fig. S6D-E and Fig. 6C-D). This points to the functional relevance of these events and distinguishes them, e.g., from the less conserved “cryptic” exons identified by examining transcriptome-wide effects of PTBP1/PTBP2 knockdown [58].

Our data suggest that PTBP1-controlled events are highly enriched for NS-CEs residing in neurodevelopmentally downregulated genes, and that its NR-CE-containing targets upregulated in neurons (Ptbp2, Gabbr1 and Dlg4; [28–31]) are more of an exception rather than the rule (Fig. S9). Interestingly, PTBP1-controlled NS-CEs are strongly enriched for the RHO GTPase effectors and actin cytoskeleton organization functions (Fig. S7B and Table S4). Another category over-represented in this set of genes relates to programmed cell death. These functions are known to play important roles in brain development and disease [65–68].

Conversely, neurodevelopmentally downregulated genes with PTBP1-independent NS-CEs are enriched for functions related to pre-mRNA splicing and other aspects of cellular RNA metabolism (Fig. S7C and Table S4). This is surprising, considering the previously established role of conserved NS-CEs in maintaining RNA-binding protein homeostasis through negative feedback loops [18, 20]. Understanding the mechanisms that allow such exons to downregulate RNA-associated factors in developing neurons is an intriguing question for future studies. It is possible that cross-regulation between functionally related proteins or developmental changes in protein modification or cellular localization patterns contribute to this process [19, 69].

An integral aspect of our work is the quantification of the extent to which NS-CEs modulate gene expression in developing neurons. Biallelic inactivation of the PTBP1-repressed NS-CEs in *Fmnl3*, *Iqgap1* and *Ripk1* suggested that these *cis*-elements facilitate downregulation of their host genes during neuronal differentiation (Fig. 5). Indeed, *Fmnl3*, *Iqgap1* and *Ripk1* lacking functional NS-CEs were expressed 4-6-fold higher compared to the wild type on differentiation days 4-6. It is also evident that NS-CEs do not function alone since some downregulation is observed even in their absence. Notably, the NMD-dependent and -independent downregulation mechanisms appear to be coordinated in time with a ±1-day precision (Fig. 5).

Further studies will be required to understand the nature of the NMD-independent repression mechanism(s) partnered with NS-CEs. We assumed transcriptional repression in our model (Fig. S12). However, the wild-type and mutant trajectories would be expected to diverge in a similar manner if AS-NMD cooperated with NMD-independent post-transcriptional repression e.g. dampening mRNAs abundance via microRNAs [70], RNA binding proteins [71], or nuclear retention of incompletely spliced mRNAs [72].

## Conclusions

We systematically elucidated the role of AS-NMD during neuronal differentiation using the newly developed bioinformatics suite factR2 and relevant experimental approraches. This revealed that NMD-stimulating cassette exons play a widespread role in promoting the downregulation of non-neuronal genes in neurons. With the increasing availability of RNA sequencing data, particularly from single-cell and long-read studies, we anticipate that the systematic use of factR2 as the transcriptome annotation tool will unveil the global impacts of AS-NMD in a variety of biological contexts.

## Materials and Methods

### DNA constructs

Plasmids p2lox and pX330-U6-Chimeric_BB-CBh-hSpCas9 were kindly provided by Michael Kyba (Addgene plasmid #34635; [46]) and Feng Zhang (Addgene plasmid #42230; [73]), respectively. pEGFP-N3 was from Clontech and pGEM-T Easy was from Promega (cat# A137A). New constructs were generated as described in Table S6 using routine molecular cloning techniques and enzymes from New England Biolabs. We used a Quikchange site-directed mutagenesis protocol with the KAPA HiFi DNA polymerase (Kapa Biosystems, cat# KK2101) to modify the *Fmnl3* minigene plasmid. Primers used for cloning, mutagenesis, and other purposes are listed in Table S7. All constructs were verified by Sanger sequencing.

### ESC culture

A2lox [46], IB10 [74], and TRE-Ngn2 (this study) mouse ESCs were cultured in a humidified incubator at 37°C, 5% CO_2_, in tissue-culture plates or dishes coated with gelatin (Millipore, cat# ES-006-B). The 2i+LIF medium [75] used for ESC maintenance contained a 1:1 mixture of Neurobasal (Thermo Fisher Scientific, cat# 21103049) and DMEM/F12 (Sigma, cat# D6421) media supplemented with 100 units/ml PenStrep (Thermo Fisher Scientific, cat# 15140122), 1 µM PD03259010 (Cambridge Bioscience, cat# SM26-2), 3 µM CHIR99021 (Cambridge Bioscience, cat# SM13-1), 0.5 mM L-Glutamine (Thermo Fisher Scientific, cat# 25030024), 0.1 mM β-mercaptoethanol (Sigma, cat# M3148), 1000 units/ml ESGRO LIF (Millipore, cat# ESG1107), 0.5× B-27 supplement without vitamin A (Thermo Fisher Scientific, cat# 12587010) and 0.5× N2 supplement. N2 100× stock was prepared using DMEM/F12 medium as a base and contained 5 mg/ml BSA (Thermo Fisher Scientific, 15260037), 2 µg/ml progesterone (Sigma, P8783-1G), 1.6 mg/ml putrescine (Sigma, P5780), 3 µM sodium selenite solution (Sigma, S5261), 10 mg/ml apo-transferrin (Sigma, T1147), and 1 mg/ml insulin (Sigma, I0516) and stored in single-use aliquots at -80°C. Cells were typically passaged every 2-3 days by treating the cultures with 0.05% Trypsin-EDTA (Thermo Fisher Scientific, cat# 15400054) for 8-10 min at 37°C. After quenching trypsin with FBS (Thermo Fisher Scientific, cat# SH30070.03E), cells were washed once with Neurobasal medium, spun down at 260×g for 4 min and plated at 1:3 to 1:6 dilution.

### Production of Dox-inducible Ngn2 knock-in ESCs

A2lox cells were pre-treated overnight with 2 µg/ml doxycycline (Dox; Sigma, cat# D9891) to activate Cre expression, trypsinized, FBS-quenched, and transfected in suspension with 1 µg of p2Lox-based plasmid pML156 (Table S6) mixed with 3 µl of Lipofectamine 2000 (Thermo Fisher Scientific, cat# 11668019) and 100 µl of Opti-MEM I (Thermo Fisher Scientific, cat# 31985070). The transfection was done in 4 ml of 2i+LIF medium at ∼2.5×10^5^ cells/ml in 6-cm bacterial dishes. Cells were pelleted 2 hours post transfection, resuspended in 2 ml of fresh 2i+LIF medium, serially diluted, and replated into a gelatinized tissue-culture 6-well plate. 350 µg/ml of G418/geneticin (Sigma, cat# 10131019) was added 36 hours post transfection and the incubation was continued for an additional 10 days with regular medium changes to allow G418-resistant cells to form colonies. These were picked, expanded, and analyzed for Dox-inducible expression of *Ngn2* by reverse-transcriptase quantitative PCR (RT-qPCR).

### Genetic inactivation of NMD-stimulating cassette exons

NS-CEs were knocked out by co-transfecting single-cell suspensions of TRE-Ngn2 cells with four plasmids encoding Cas9 and distinct gene-specific CRISPR gRNAs, as well as the pEM584 plasmid [76] encoding a blasticidin resistance gene. 1.2 µg of a mixture containing equal amounts of each of these plasmids was combined with 3 µl of Lipofectamine 2000 and 100 µl of Opti-MEM I. TRE-Ngn2 cells were transfected with the mixture in 4 ml of 2i+LIF at ∼2.5×10^5^ cells/ml in 6-cm bacterial dishes. Cells were collected 2 hours post transfection, spun down, and the cell pellets were serially diluted in fresh 2i+LIF prior to plating in 6-well format. Antibiotic selection with 8 µg/ml blasticidin (Sigma, cat# 15205) was started 24 hours post transfection and lasted for 3 days. Cells were then cultured in 2i+LIF medium for another 6-10 days. Surviving colonies were picked, expanded for 3-4 days, and PCR-genotyped (see below). In cases where heterogeneity was suspected, the clones were dissociated to single-cell suspensions and allowed to form colonies (without blasticidin). These were picked, expanded, genotyped, and used for subsequent analyses.

### Genotyping

Genomic DNA was extracted and analyzed using the PCRBIO Rapid Extract PCR Kit (PCR Biosystems; cat# PB10.24-08) following the manufacturer’s protocol. Amplified DNA fragments were resolved by electrophoresis in 1% agarose gels alongside GeneRuler 1 kb Plus DNA Ladder (Thermo Fisher Scientific, cat# SM1331). CRISPR/Cas9-induced mutations in AS-NMD-regulated genes were characterized with primers designed to amplify either short (∼0.5 kb) or long (∼2 kb) fragments centered on the targeted exon. Gel-purified PCR fragments were cloned into the pGEM-T Easy vector (Promega, cat# A137A) and sequenced using the Sanger’s method.

### Transient transfection

Cells were counted using a hemocytometer, and ∼2×10^5^ cells in 1 ml of 2i+LIF medium were seeded per well of a gelatinized 12-well plate and immediately transfected with 50 pmol of an appropriate siRNA (Horizon Discovery/Dharmacon; Table S8) premixed with 2.5 µl of Lipofectamine 2000 and 100 µl of Opti-MEM I, as recommended. In experiments involving simultaneous knockdown of two genes, 100 pmol of mixed siRNAs was combined with 4 µl of Lipofectamine 2000 and 100 µl Opti-MEM I. Cells were exposed to siRNA-containing complexes overnight, followed by a medium change to fresh 2i+LIF next morning. RNAs were extracted 48 hours post transfection. In experiments requiring CHX treatment, cells grown for 48 hours following siRNA transfection were supplemented with 100 µg/ml CHX (Sigma, cat# C4859) or an equal volume of DMSO (Sigma, cat# D2650) and incubated for an additional 6 hours prior to RNA extraction. In minigene experiments, cells were transfected with 1 µg of a corresponding plasmid mixed with 3 µl of Lipofectamine 2000 and 100 µl of Opti-MEM I and incubated for 24 hours. Alternatively, cells were pre-treated with siRNAs for 24 hours and then transfected with 1 µg of minigene plasmid, as above, and incubated for another 24 hours prior to RNA extraction.

### Inducible neuronal differentiation

We have adapted previously published protocols for *Ngn2*-induced glutamatergic neuronal differentiation [44, 45] (Table S9). On the day of *Ngn2* induction (day 0), cell culture plates (12-well plates were Corning™ Costar™ cat# 3513 and 6-well plates were Thermo Fisher Scientific Nunc™ cat# 140675) were coated with Geltrex^TM^ (Gibco, cat# A1413302), diluted 1:50 in cold DMED/F12, for 2 hours in a humidified incubator at 37°C. Geltrex was aspirated and TRE-Ngn2 ESCs were immediately plated into wells, without washing, in the iN-D0 medium containing a 1:1 mixture of Neurobasal (Thermo Fisher Scientific, cat# 21103049) and DMEM/F12 (Sigma, cat# D6421) supplemented with 100 units/ml PenStrep (Thermo Fisher Scientific, cat# 15140122), 1×N2 (Gibco, cat# 17502048), 1×B27 with retinoic acid (Gibco, cat# 17504044), 1 µg/ml laminin (Sigma, cat# L2020), 20 µg/ml insulin (Sigma, cat# I0516), 500 µM L-glutamine (Gibco, cat# 25030-024), 2 mM db-cAMP (Sigma, cat# D0627), 10 ng/ml NT-3 (Miltenyi Biotec, cat# 130-093-973), 100 µM β-mercaptoethanol and 2 µg/ml Dox. If not stated otherwise, we seeded 2×10^4^ ESCs per well of a Geltrex-coated 12-well plate for subsequent RT-PCR and RT-qPCR analyses and 1-3×10^3^ cells per well of a 12-well plate containing a Geltrex-coated 18-mm glass coverslip (VWR, cat# 631-1580) for immunostaining. The cells attached to the wells were then cultured in a humidified incubator at 37°C, 5% CO_2_.

On day 2, the medium was changed completely to the iN-D2 medium, similar to iN-D0 except containing 200 µM L-ascorbic acid (Sigma, A4403) instead of β-mercaptoethanol and 1 µg/ml instead of 2 µg/ml Dox. Starting from day 4, we replaced only ∼50% of conditioned medium with fresh Dox-free medium. The iN-D4 medium used for this purpose on day 4 had the same composition as iN-D2 but lacked Dox. In short-term differentiation experiments, samples were collected on day 6 without further medium changes. In experiments where induced neurons were maintained longer than 6 days, the half-volume changes on day 6 onwards were done using the iN-D6 medium containing Neurobasal supplemented with 1×B27 with retinoic acid, 1×CultureOne supplement (Gibco, cat# 15674028), 500 µM L-glutamine, 1 µg/ml laminin, 100 units/ml PenStrep, 200 µM L-ascorbic acid, 10 ng/ml NT-3, 10 ng/ml BDNF (Miltenyi Biotec, cat# 130-093-811), and 2 mM db-cAMP.

### RT-qPCR and RT-PCR analyses

Total RNAs were purified using the EZ-10 DNAaway RNA Miniprep Kit (BioBasic, cat# BS88136), as recommended. Reverse transcription (RT) was performed at 50°C for 40 min using SuperScript IV reagents (Thermo Fisher Scientific, cat# 18090200), 5 µM of random decamer (N10) primers, and 2 units/µl of murine RNase inhibitor (New England Biolabs, M0314L). In minigene experiments, purified RNAs were additionally incubated with 2 units of RQ1-DNAse (Promega, cat# M6101) per 1 µg of RNA at 37°C for 30 min to eliminate plasmid DNA contamination. RQ1-DNAse was inactivated by adding the Stop Solution. The RNAs were then immediately reverse-transcribed as described above.

cDNA samples were analyzed by qPCR using a Light Cycler®96 Real-Time PCR System (Roche) and the qPCR BIO SyGreen Master Mix (PCR Biosystems; cat# PB20.16). All RT-(q)PCR primers are listed in Table S7. RT-qPCR signals were normalized to expression levels of housekeeping genes selected based on our RNA-seq data. *Cnot4* was used for longitudinal differential expression analyses. *Gars* was used to compare between CHX- and DMSO-treated samples.

To analyze NS-CE inclusion, cDNA samples were amplified using DreamTaq DNA Polymerase (Thermo Fisher Scientific cat# EP0702) or ROCHE Taq DNA polymerase (Sigma-Aldrich, cat# 11596594001). The RT-PCR products were then resolved by electrophoresis in 2% agarose gels alongside GeneRuler 1 kb Plus DNA Ladder. Band intensities were quantified using Image Studio Lite (Version 5.2) (LI-COR Biosciences). Percent spliced in (PSI) values were calculated by normalizing the intensity of the NS-CE inclusion product by combined intensity of the inclusion and skipping products and multiplying the quotient by 100.

### TRE-Ngn2 samples for RNA-sequencing

For RNA-Seq, TRE-Ngn2 cells were differentiated into neurons in a 6-well plate format as described above. We plated 6-8×10^4^ cells per well to produce day 3, 6, 12 and 24 differentiated samples. Day 0 non-induced samples were generated by plating TRE-Ngn2 ESCs in 2i+LIF at 1×10^5^ per well and then lysing the culture 3 days later. All time points were lysed using TRIzol reagent (Thermo Fisher Scientific, cat# 15596026). The lysates were immediately frozen, stored at -80°C, and processed simultaneously using the TRIzol Plus RNA Purification Kit (Thermo Fisher Scientific cat# 12183555). RNAs were eluted in nuclease-free water (Invitrogen, cat# AM9939).

### Natural samples for RNA sequencing

To identify natural AS-NMD targets, cultures of wild-type (IB10) ESCs, NPC neurospheres, and primary neurons were treated with either DMSO or 100 µg/ml CHX for 6 h at 37°C. Alternatively, we prepared nuclear and cytoplasmic fractions from untreated cells using a PARIS™ kit (Thermo Fisher Scientific, cat# AM1921).

Primary NPC cultures were obtained by dissecting embryonic day 14.5 (E14.5) mouse embryonic cortices in Hank’s Balanced Salt Solution (1×HBSS, Thermo Fisher Scientific; cat# 14025092) and dissociating them by trituration. The NPCs were then cultured as neurospheres in reconstituted NeuroCult® Proliferation Kit medium (STEMCELL Technologies, cat# 05702) supplemented with 20 ng/ml recombinant human EGF (STEMCELL Technologies, cat# 78006). For passaging, neurospheres were dissociated with NeuroCult® Chemical Dissociation Kit (STEMCELL Technologies, cat# 05707), as recommended. We also prepared adherent NPC cultures for immunofluorescence experiments by plating dissociated neurospheres to polyornithine (Sigma; cat# P3655-10MG) and fibronectin (STEMCELL Technologies, cat# 07159) coated dishes.

Primary cortical neurons were isolated from E15.5 mouse embryos and cultured as described [77]. Briefly, cortices were dissociated with 2.5% trypsin (Thermo Fisher Scientific, cat# 15090046) and plated on poly-L-lysine (Sigma, cat# P2636) coated 6-well plates or 18-mm round coverslips in Minimum Essential Media (MEM) with L-glutamine (Thermo Fisher Scientific, cat# 11095080), 0.6% glucose (Fisher Scientific, cat# 10335850) and 10% horse serum (GIBCO/Life Technologies) at 1×10^6^ neurons per well and 2.5×10^4^ neurons per coverslip.

Neurons in 6-well plates were cultured for 5 days without glial feeders, whereas neurons attached to coverslips were transferred to wells containing a monolayer of newborn rat astrocytes [77], and cultured in neuronal maintenance medium (MEM with L-glutamine, 0.6% glucose and 1× Neurocult SM1 neuronal supplement (STEMCELL Technologies, cat# 05711)) for 21 days replacing half of the medium every 3-4 days.

Astrocytes for long-term neuronal cultures were prepared by dissociating newborn rat cortices with 2.5% trypsin and 1 mg/ml DNase (Merck, cat# 10104159001) and plating the cells in MEM with L-glutamine supplemented with 0.6% glucose, 10% FBS (GE Hyclone, cat# HYC85), 100 units/ml penicillin and 100 µg/ml streptomycin (Thermo Fisher Scientific, cat# 15140122) at 2×10^6^ cells per 25-cm^2^ flask [77]. Medium was changed once after 3 days, and the cultures were maintained for a total of 7 days to allow astrocytes to expand to ∼90% confluence.

We isolated RNAs from the whole-cell and fractionated material using either the Purelink™ RNA mini kit (Thermo Fisher Scientific, cat# 12183025; IB10 ESCs and NPCs) or the RNeasy micro kit (Qiagen, cat# 74004; primary neurons).

### RNA sequencing

Total RNA concentration was estimated using Nanodrop and measured with the Qubit RNA BR Assay Kit (Fisher Scientific cat# Q10211). Additionally, RNA quality was evaluated using the Agilent Technologies RNA ScreenTape Assay (5067-5576 and 5067-5577).

For the TRE-Ngn2 time series, intact poly(A) RNA was purified from total RNA samples (100-500 ng) with oligo(dT) magnetic beads. Stranded mRNA sequencing libraries were then prepared as described using the Illumina TruSeq Stranded mRNA Library Prep kit (20020595) and TruSeq RNA UD Indexes (20022371). Purified libraries were qualified on an Agilent Technologies 2200 TapeStation using a D1000 ScreenTape assay (cat# 5067-5582 and 5067-5583). The molarity of adapter-modified molecules was defined by quantitative PCR using the Kapa Biosystems Kapa Library Quant Kit (cat# KK4824). Individual libraries were normalized to 1.30 nM in preparation for Illumina sequence analysis. Sequencing libraries were chemically denatured and applied to an Illumina NovaSeq flow cell using the NovaSeq XP chemistry workflow (20021664). Following transfer of the flowcell to an Illumina NovaSeq instrument, a 2×51 cycle paired end sequence run was performed using the NovaSeq S1 reagent Kit (20027465).

For the natural samples, intact poly(A) RNA was purified from total RNA (100-500 ng) with oligo(dT) magnetic beads and stranded mRNA sequencing libraries were prepared as described using the Illumina TruSeq Stranded mRNA Library Preparation Kit (RS-122-2101, RS-122-2102). Purified libraries were qualified on an Agilent Technologies 2200 TapeStation using a D1000 ScreenTape assay (cat# 5067-5582 and cat# 5067-5583). The molarity of adapter-modified molecules was defined by quantitative PCR using the Kapa Biosystems Kapa Library Quant Kit (cat# KK4824). Individual libraries were normalized to 5 nM and equal volumes were pooled in preparation for Illumina sequence analysis. Sequencing libraries (25 pM) were chemically denatured and applied to an Illumina HiSeq v4 single read flow cell using an Illumina cBot. Hybridized molecules were clonally amplified and annealed to sequencing primers with reagents from an Illumina HiSeq SR Cluster Kit v4-cBot (GD-401-4001). The flowcell was transferred to an Illumina HiSeq 2500 instrument (HCSv2.2.38 and RTA v1.18.61) and a 50-cycle single-read sequence run was performed using HiSeq SBS Kit v4 sequencing reagents (FC-401-4002). The library preparation and sequencing steps were performed by the High-Throughput Genomics facility at the Huntsman Cancer Institute, University of Utah, USA.

### Immunofluorescence analyses

Non-induced TRE-Ngn2 ESCs were plated in 2i+LIF medium onto 18-mm gelatin-coated coverslips (VWR, cat# 631-1580) at 6×10^4^ cells per well of a 12-well plate. Dox-induced TRE-Ngn2 cells were plated onto 18 mm coverslips coated with Geltrex, as described above. Cells were fixed with 4% PFA in 1×PBS for 15 min at room temperature. After 3 washes with 1×PBS, cells were permeabilized with 0.1% Triton X-100 in 1×PBS for 5 min, followed by another 3 washes with 1×PBS. The coverslips were then incubated in the blocking buffer (5% horse serum, 5% goat serum, 1% BSA in 1×PBS) for 1 hour at room temperature. Blocked samples were incubated with primary antibodies (Table S10) diluted in the blocking buffer at 4°C in a humidified chamber overnight. Next day, coverslips were washed 3 times with 1×PBS and incubated with Alexa Fluor-conjugated secondary antibodies diluted 1:400 in the blocking buffer at room temperature for 1 hour. Coverslips were then washed 3 times with 1×PBS and stained with 0.5 µg/ml DAPI (Thermo Fisher Scientific, cat# D1306) in 1×PBS for 5 min. Coverslips were mounted on microscope slides in a drop of ProLong Gold Antifade Mountant solution (Thermo Fisher Scientific, cat# P36934). Images were taken using a ZEISS Axio Observer Z1 Inverted Microscope with alpha Plan-Apochromat 100x/1.46 oil immersion objective.

### Bioinformatics

RNA-seq data were quality-controlled by FastQC (http://www.bioinformatics.babraham.ac.uk/projects/fastqc/) and trimmed by Trimmomatic [78]. HISAT2 [50] index and splice site files were prepared using the GRCm39 mouse genome (GRCm39.primary_assembly.genome.fa) and M26 transcriptome (gencode.vM26.primary_assembly.annotation.gtf) files from GENCODE (ftp://ftp.ebi.ac.uk/pub/databases/gencode/Gencode_mouse/release_M26/):

hisat2-build -p [n_threads] GRCm39.primary_assembly.genome.fa GRCm39_genome hisat2_extract_splice_sites.py gencode.vM26.primary_assembly.annotation.gtf > M26.ss.txt

We then aligned our longitudinal TRE-Ngn2 differentiation data using HISAT2: hisat2 -p [n_threads] -k 50 --fr --rna-strandness RF --dta-cufflinks --no-unal \

--known-splicesite-infile M26.ss.txt \

-x GRCm39_genome \

-1 [read_1.fastq] \

-2 [read_2.fastq] \

-S [aligned.sam]

SAM files were converted to the BAM format and optionally concatenated, sorted and indexed using SAMtools [79]. To produce CPM-normalized strand-specific RNA-seq coverage plots, sorted and indexed [aligned.bam] files were converted into the bedGraph format using deepTools [80]:

bamCoverage -p [n_threads] -bs 1 -of bedgraph \

--filterRNAstrand forward \

--effectiveGenomeSize 2654621837 --normalizeUsing CPM \

-b [aligned.bam] -o [forward.bedGraph] bamCoverage -p [n_threads] -bs 1 -of bedgraph \

--filterRNAstrand reverse \

--effectiveGenomeSize 2654621837 --normalizeUsing CPM \

-b [aligned.bam] -o [reverse.bedGraph]

The resultant [forward.bedGraph] and [reverse.bedGraph] files were then visualized in IGV [81].

To identify novel transcripts enriched for AS-NMD events, we first analyzed individual stage-specific DMSO- and CHX-treated samples by StringTie [50]:

stringtie -p [n_threads] --rf -M 1 -u -f 0.05 \

-G gencode.vM26.primary_assembly.annotation.gtf \

-o [sample_specific.gtf] \ [sample_specific.bam]

Different sample_specific.gtf files were then merged by cuffmerge [82]: cuffmerge -p [n_threads] --min-isoform-fraction 0.05 \

-g gencode.vM26.primary_assembly.annotation.gtf \

[list_of_sample_specific_gtfs_to_merge.txt]

and novel transcripts in the resultant merged.gtf file were shortlisted by removing entries with intron chains identical to those in gencode.vM26.primary_assembly.annotation.gtf (class code “=”):

grep -v -P “class_code \“=\”;” merged.gtf > novel.gtf

The remaining transcripts were analyzed by GffCompare [83] to allocate them to known genes where possible:

gffcompare novel.gtf -r gencode.vM26.primary_assembly.annotation.gtf

We then used R [84] and factR2 (https://github.com/f-hamidlab/factR2) to annotate the resultant gffcmp.merged.gtf file, combine it with the gencode.vM26.primary_assembly.annotation.gtf reference and extract AS-NMD events from thus produced extended_M26.gtf:

library(factR2) library(rtracklayer)

custom.gtf <-import(“gffcmp.merged.gtf”)

ref.gtf <- import(“gencode.vM26.primary_assembly.annotation.gtf”) seqlevels(ref.gtf) <- unique(c(seqlevels(custom.gtf), seqlevels(ref.gtf))) extended_M26.gtf <- c(ref.gtf, custom.gtf)

factRobj <- createfactRObject (extended_M26.gtf, “vM26”) factRobj <- buildCDS(factRobj)

factRobj <- predictNMD(factRobj) factRobj <- testASNMDevents(factRobj)

factRobj <- getAScons(factRobj, type=“flanks”, padding=100)

exportGTF(factRobj, “extendend_M26.gtf”)

exportTable(factRobj, “extended_M26_NMD_exons.txt”, “exons”)

Transcript abundance in our TRE-Ngn2 differentiation experiment was quantified using Kallisto [85] with an extended_M26.gtf-based index:

kallisto quant -t [n_threads] --rf-stranded \

-i extended_M26.index -o [out_dir] \ [read_1.fastq] [read_2.fastq]

Kallisto-deduced transcript-specific counts for individual biological replicates were imported into R using the tximport package [86] and analyzed by DESeq2 [52]. Genes not represented by ≥5 sequencing reads in ≥2 samples were considered poorly expressed and excluded from subsequent analyses. DMSO and CHX data for specific differentiation time points were compared pairwise using the Wald test:

dds <- DESeq(dds, fitType=’local’)

Differentially expressed genes in DMSO time-course data were analyzed by the likelihood-ratio test:

dds_lrt <-DESeq(dds_lrt, test=“LRT”, reduced = ∼ 1)

Unless indicated otherwise, genes with FDR (calculated as Benjamini-Hochberg-adjusted *P*-value) <0.001 were considered differentially regulated. The PCA plot in Fig. S2A was produced by DESeq2’s plotPCA function using VST-transformed expression values (𝑉𝑆𝑇_*gene*_), which were calculated for all detectably expressed genes by the vst(dds, blind=FALSE) function of DESeq2. 𝑉𝑆𝑇_*gene*_values were also used for correlation and developmental trend analyses. Heatmaps in Fig. S2B and Fig. S9A were produced using the ComplexHeatmap R package and centered and scaled 𝑉𝑆𝑇_*gene*_values for manually selected marker genes.

Alternative splicing patterns were analyzed by Whippet with an extended_M26.gtf-based index:

julia whippet-quant.jl [read_1.fastq] [read_2.fastq] --biascorrect \

-x extended_M26.whippet.graph.jls -o [out_dir]

Condition-specific psi.gz files produced by the above code were then compared for each differentiation stage as follows:

julia whippet-delta.jl -s 1 -a condition_A.psi.gz -b condition_B.psi.gz \

-o [output_prefix]

The difference (Δ) between condition-specific percent spliced in values (PSI; defined in Whippet as a fraction rather than %) for CHX-treatment and nucleus-cytoplasm fractionation experiments was calculated as:

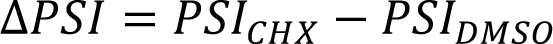

and

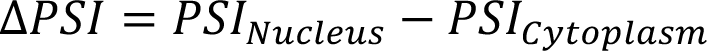

respectively.

We used the following filters to shortlist regulated NMD-stimulating (NS) and NMD-repressing (NS) events. NS events, ΔPSI>0.1 and probability>0.9; NR events, ΔPSI<-0.1 and probability>0.9. To retain events that control mRNA sensitivity to NMD but are nearly completely included or excluded at the splicing level, we also shortlisted NS events where PSI averaged across conditions was >0.9, and NR events where the average was <0.1. Shortlisted events from the cassette/core exon (CE), alternative donor/5’ splice site (AD), alternative acceptor/3’ splice site (AA), and retained intron (RI) groups were used for further analyses.

For correlation and trend analyses we used facR2’s function testGeneCorr subjecting PSI values to arcsine-square root variance-stabilizing transformation [87] and adding the 2/π coefficient to maintain transformed PSI within the [0, 1] interval:

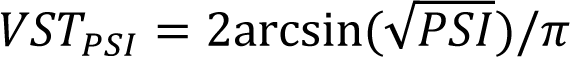

Correlation between AS-NMD splicing pattern and gene expression was then calculated as Pearson’s product-moment correlation between 𝑉𝑆𝑇_*gene*_and 𝑉𝑆𝑇_*PSI*_ for NMD-repressing events and between 𝑉𝑆𝑇_*gene*_ and 1 − 𝑉𝑆𝑇_*PSI*_ for NMD-stimulating events.

Evolutionary conservation of the intronic sequence context of AS-NMD events was analyzed using factR2’s function getASCons and the mm39 PhastCons 35-vertebrate conservation track from the UCSC genome browser (http://hgdownload.soe.ucsc.edu/goldenPath/mm39/phastCons35way/). For each event type, we examined a 200-nt sequence, defining the context as 100-nt intronic sequences preceding and following CEs, the first and last 100 nt of RIs, 200-nt intronic sequences following ADs, and 200-nt intronic sequences preceding AAs.

PTBP1-regulated events were shortlisted by Whippet analyses of ESCs and NPCs, where PTBP1 was knocked down alone or together with its functionally similar paralog, PTBP2 (|ΔPSI|>0.1; probability>0.9 cutoffs). If the PSI values changed significantly in response to multiple treatments, individual PTBP1 knockdown experiments were given priority over combined PTBP1 and PTBP2 knockdowns. Moreover, ESCs were prioritized over NPCs. We additionally requested that the event’s PSI correlates with *Ptbp1* expression levels (Pearson’s *P*<0.05) in our TRE-Ngn2 RNA-seq differentiation dataset and that the sign of the correlation coefficient (Pearson’s *r*) is opposite to the sign of ΔPSI in response to PTBP1 knockdown in ESCs or NPCs.

We estimated the proportions of different cell types in differentiating TRE-Ngn2 cultures using the R package MuSiC [48]. Raw gene expression counts from our bulk RNA-seq dataset were compared with mouse single-cell RNA-seq data for ESCs (GSM2098554), NPCs (GSE67833; [88]), and adult cortices (GSE185862; [89]). We first converted count matrices to the ExpressionSet object format:

library(Biobase)

metadata <- data.frame(row.names = colnames(counts), group = colnames(counts)) metalabels <- data.frame(labelDescription=c(“group”))

metadata <- AnnotatedDataFrame(data=metadata, varMetadata=metalabels) bulk.eset <- ExpressionSet(counts, phenoData = metadata)

and then performed cellular deconvolution using MuSiC’s music_prop function. The output data frame was processed and visualized in ggplot2 [90].

Genes were clustered according to their neurodevelopmental expression trajectories using DP_GP_cluster [53] as follows:

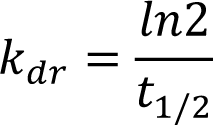

DP_GP_cluster.py -i [expression.txt] -o [output_prefix] \

--fast -n 1000 --max_iters 1000 \

--check_burnin_convergence --check_convergence \

--cluster_uncertainty_estimate --plot -p pdf

where expression.txt was a table with gene IDs and averages of 𝑉𝑆𝑇_*gene*_values for each differentiation time point. The optimal_clustering.txt output file was used for further analyses.

To estimate AS-NMD contributions to gene downregulation in development, we modeled changes in mRNA abundance as a function of time (𝑅(𝑡)) depending on mRNA synthesis rate (𝑣_*sr*_) and the decay constant (𝑘_*dr*_) related to mRNA half-life (𝑡_1/2_) [91, 92] as follows:

For simplicity, we assumed that the rate at which productively spliced mRNA is synthesized can decrease through the onset of transcriptional repression (𝑟_*t*_) or/and an increase in AS-NMD repression (𝑟_*n*_) at specific time points within the day 2-4 period, whereas the 𝑘_*dr*_ remains constant. We also assumed that NMD-sensitive isoforms are much less stable than productively spliced mRNA, and therefore, ignored their contribution to 𝑅(𝑡). This produced the following equation, which was solved using the ode45 function or the R package pracma:

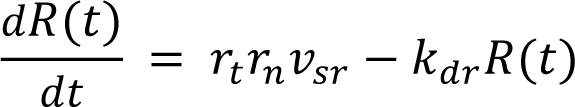

In the model described in Fig. S12, we used published transcriptome-wide estimates for 𝑣_*sr*_(1.76 mRNA molecules per cell per hour; the median rate measured for mouse NIH 3T3 cells; [92]) and 𝑡_1/2_ (7.08 hours; the median half life for proliferating and differentiating mouse ESCs; [93]). However, similar results were obtained when we estimated the 𝑣_*sr*_ and 𝑡_1/2_ parameters by averaging the *Fmnl3-*, *Iqgap1-*, and *Ripk1-*specific values (𝑣_*sr*_=2.63 mRNA molecules per cell per hour in NIH 3T3 cells and 𝑡_1/2_=5.11 hours in proliferating (LIF+) and differentiating (RA+) mouse ESCs). 𝑟_*t*_ was set to 1 in early development. To estimate 𝑟_*t*_ following the onset of repression in developing neurons, we averaged the ratios between expression levels on days 5-6 and days 0-1 for the *Fmnl3*, *Iqgap1*, and *Ripk1* mutants in Fig. 5A-C. This resulted in 𝑟_*t*_ = 0.39. We estimated the initial (stem cell-specific) and the final (neuron-specific) 𝑟_*n*_ values from the 𝑃𝑆𝐼_*CHX*_ data (Fig. 4B):

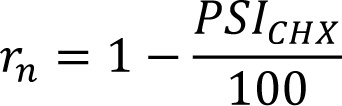

by averaging day-0-1 and day-5-6 *Fmnl3*, *Iqgap1* and *Ripk1* 𝑃𝑆𝐼_*CHX*_ values, respectively. This yielded the initial 𝑟_*n*_ = 0.78 and the final 𝑟_*n*_ = 0.12.

Single-cell RNA-sequencing data from developing mouse dentate gyrus were downloaded from https://www.ncbi.nlm.nih.gov/geo/query/acc.cgi?acc=GSE95753. A total of 2000 FASTQ files from the granule cell lineage were sampled and combined into four groups to represent the transcriptomes of the four distinct cell subpopulations: radial glia-like cells, neuronal intermediate progenitors, neuroblasts, and granule cells. Pooled FASTQ files were aligned to the mm39 genome by HISAT2, and custom transcriptomes were constructed using StringTie2, as described above. Group-specific GTF files were then merged:

stringtie --merge -F 0.5 -T 0.5 -G gencode.vM26.primary_assembly.annotation.gtf –o stringtie.merged.gtf [*.gtf]

The stringtie.merged.gtf file was processed by factR2 to annotate AS-NMD events and score the interspecies conservation of intronic regions flanking NS-CEs. Transcript abundances and splicing patterns in individual cells were analyzed by Kallisto and Whippet, respectively, with stringtie.merged.gtf-based indexes. In this analysis, high-quality NS-CEs were identified as events predicted by factR2 to stimulate NMD and showing a correlation (Pearson’s *r*>0.2 and *P*<0.05) between NMD-protective splicing and gene expression after applying appropriate variance-stabilizing transformations. Cassette exons that were not predicted to be NMD-inducing by factR2 were used as a control group.

The source code and a detailed user manual for factR2 are available on GitHub (https://github.com/f-hamidlab/factR2). The key bioinformatics procedures described above are additionally available on GitHub (https://github.com/f-hamidlab/Zhuravskaya-ASNMD).

### Statistical analyses

Unless stated otherwise, all statistical procedures were performed in R. We repeated experiments at least three times and showed the results as averages ±SD. Data obtained from RT-(q)PCR quantifications were typically analyzed using a two-tailed Student’s t-test assuming unequal variances. Other types of data were compared using two-tailed Wilcoxon rank sum test, one-sided Kolmogorov-Smirnov (KS) test or Fisher’s exact test. Where necessary, p-values were adjusted for multiple testing using Benjamini-Hochberg correction (FDR). Correlation analyses were done using Pearson’s product-moment and Kendall’s rank correlation methods, as specified in the text. Numbers of experimental replicates, *P*-values and the tests used are indicated in the Figures and/or Figure legends.

## Acknowledgements

We thank Snezhka Oliferenko for valuable discussions, Brian Dalley for the help with RNA sequencing, and Michael Kyba and Feng Zhang, for reagents. This work was supported by the Biotechnology and Biological Sciences Research Council (BB/M001199/1, BB/M007103/1, and BB/V006258/1).

---

**Figure S1.**
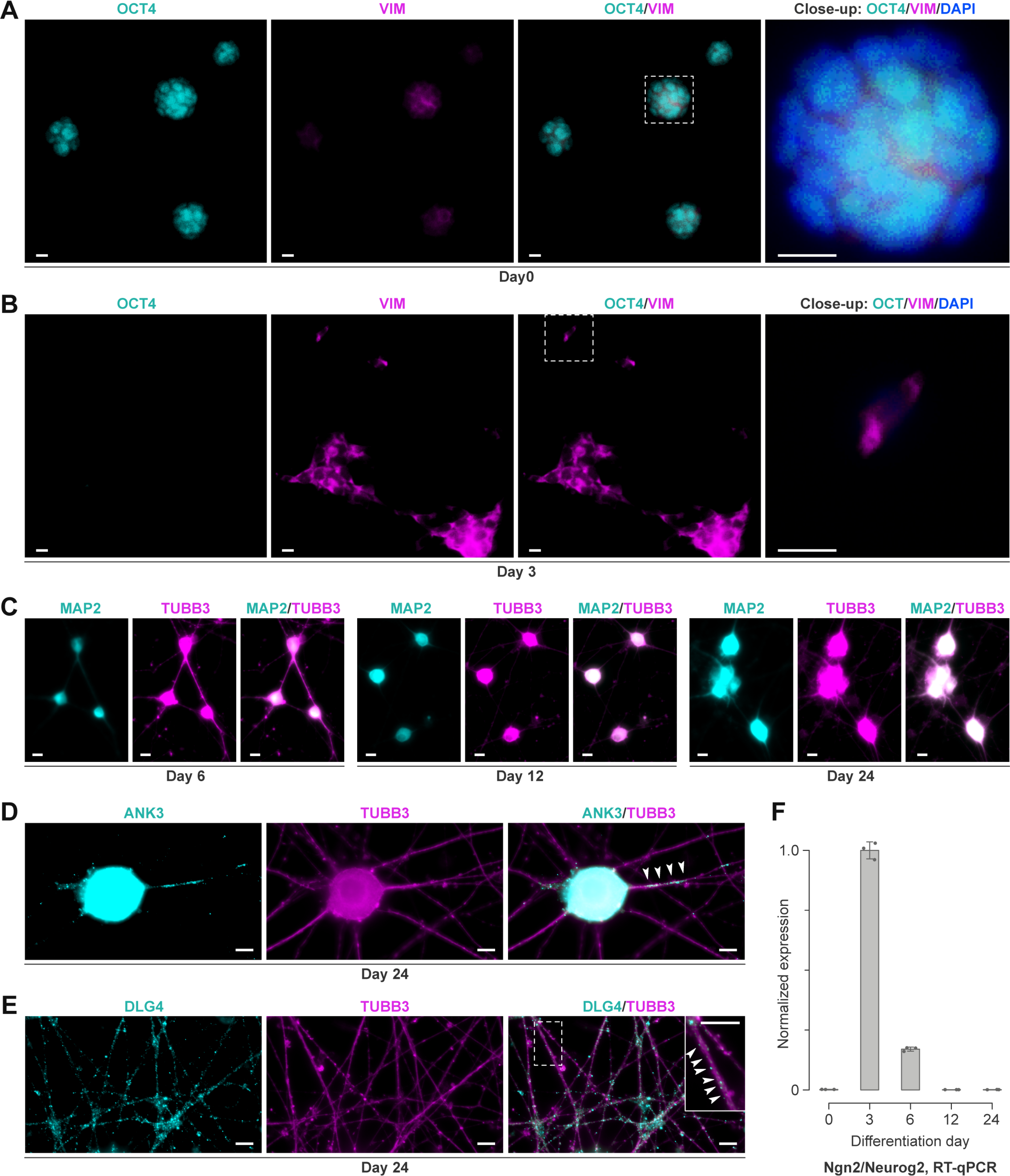
Expression of developmental markers in Dox-induced TRE-Ngn2 cultures. **(A-B)** TRE-Ngn2 cells fixed on differentiation day 0 (A) or day 3 (B) were immunostained for the ESC marker OCT4/POU5F1 (Abcam, ab19857; 1:500) and the NPC-enriched marker VIM (Abcam, ab8978; 1:500). Note that OCT4 is expressed in day-0 but not day-3 cells and that VIM is strongly upregulated on day 3. **(C)** TRE-Ngn2 samples differentiated for 6-24 days were immunostained for an early neuronal marker, TUBB3/tubulin βIII (Covance, PRB-435P; 1:5000), and a later neuronal marker, MAP2 (Covance, PCK-554P, 1:1000). Note an increase in MAP2 expression compared to the TUBB3 signal at later stages of differentiation. **(D-E)** Day-24 TRE-Ngn2 cultures were stained for mature neuronal markers: (D) ANK3/ANKG (NeuroMab, 75-146, 1:500) and (E) DLG4/PSD-95 (NeuroMab, 75-028, 1:500). Co-staining for TUBB3 was used to visualize the overall neuronal morphology. Arrowheads indicate an ANK3-positive axon initial segment and DLG4 puncta, respectively. (A, B, E) Dashed rectangles, areas magnified in the close-ups. (A-E) Scale bars, 10 µm. (**F**) RT-qPCR analysis shows that the *Ngn2*/*Neurog2* expression in differentiating TRE-Ngn2 cells peaks at the NPC stage (day 3) and declines at later stages. This mimics the natural regulation of *Ngn2* in neurodevelopment [45].

**Figure S2.**
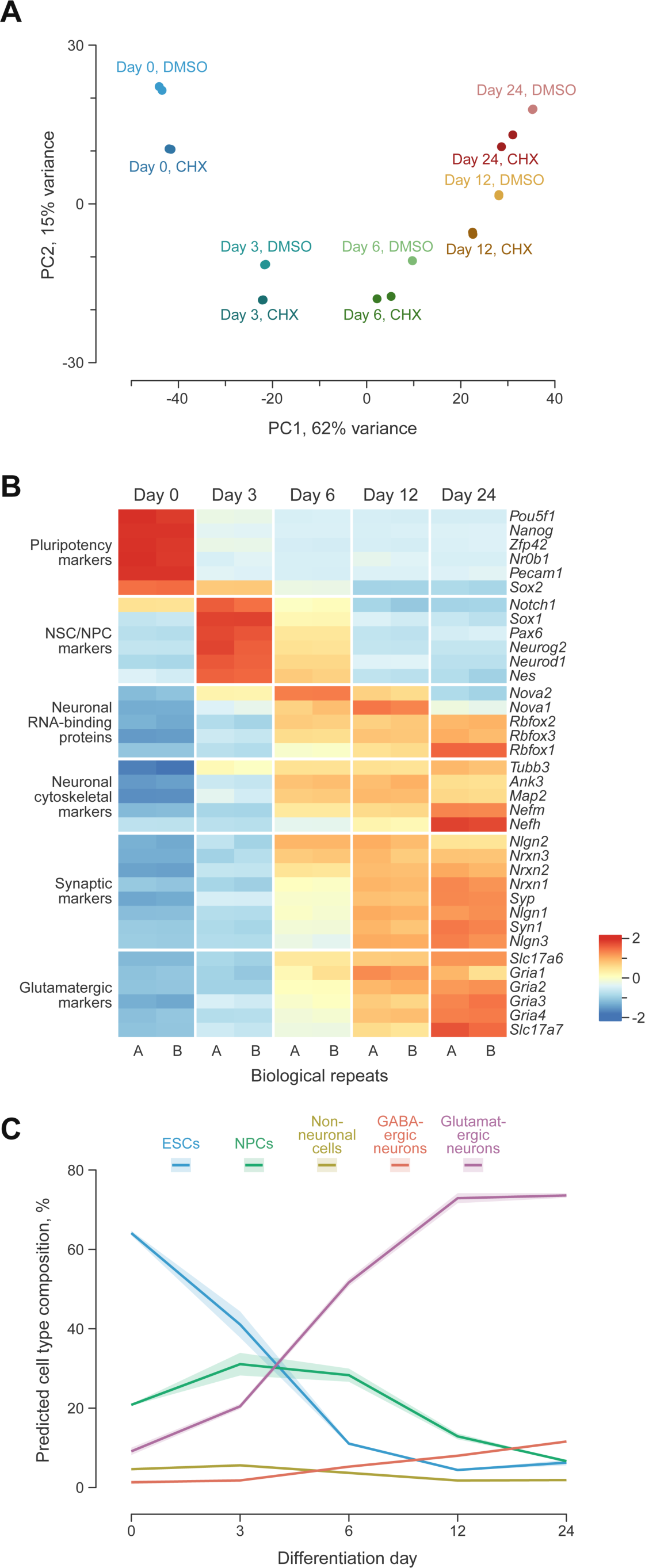
Initial characterization of the TRE-Ngn2 RNA-seq data. **(A)** Principal component analysis (PCA) showing that the transcriptome of TRE-Ngn2 cells changes as a function of development and responds to the CHX treatment. **(B)** Heat map analysis of stage-specific markers in the control-treated TRE-Ngn2 RNA-seq samples. **(C)** Changes in the cellular composition of differentiating TRE-Ngn2 cultures deconvolved by MuSiC [48].

**Figure S3.**
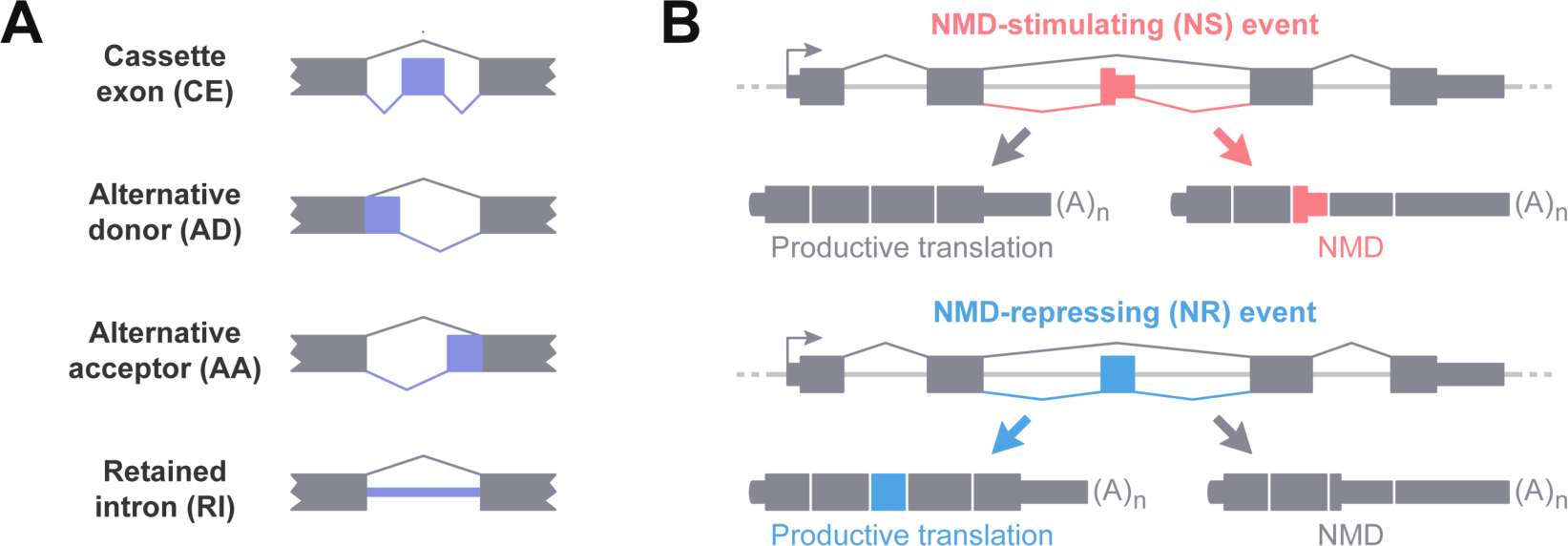
Classification of AS-NMD events in factR2. **(A)** The key types of alternatively spliced events examined by factR2. **(B)** The opposite effects of NMD-stimulating (NS) and NMD-repressing (NR) events on gene expression illustrated for cassette exons, as an example.

**Figure S4.**
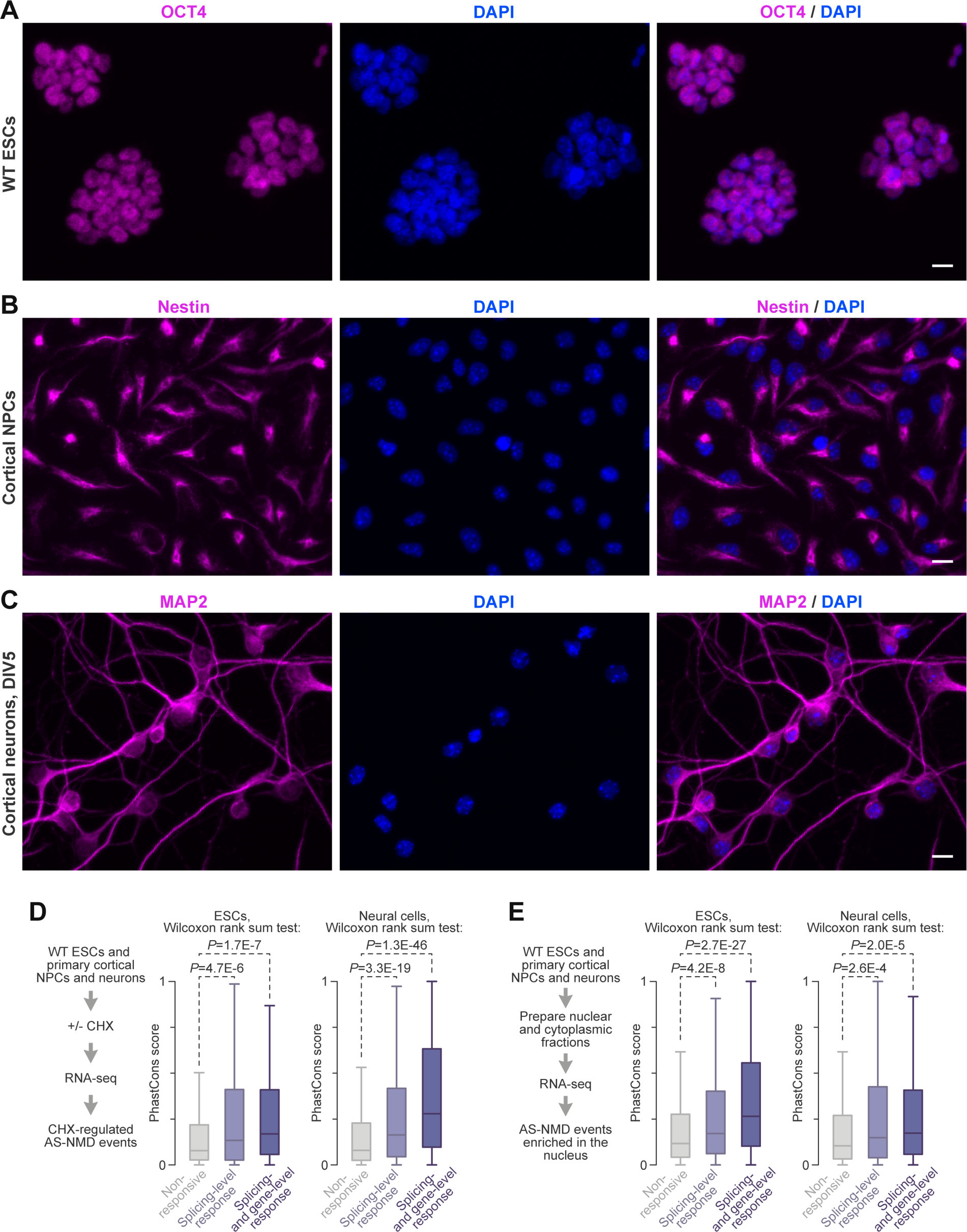
Detailed analyses of wild-type ESCs and primary neural cells. **(A-C)** Immunofluorescence analyses of (A) wild-type ESCs (WT; IB10 line) for OCT4 expression; (B) cortical NPCs for NES/Nestin expression; and (C) cortical neurons cultured 5 days in vitro for MAP2 expression. Scale bars, 10 µm. **(D)** Interspecies conservation of intronic sequence context of the AS-NMD events responding to CHX in the WT ESCs or primary neural cells and additionally present in different groups of the TRE-Ngn2 events. Higher PhastCons scores [94] indicate stronger conservation across vertebrates. See Fig. 1F for further details. **(E)** Interspecies conservation of intronic sequence context of the AS-NMD events enriched in the nucleus of the WT ESCs or primary neural cells and additionally present in different groups of the TRE-Ngn2 events. See Fig. 1G for further details.

**Figure S5.**
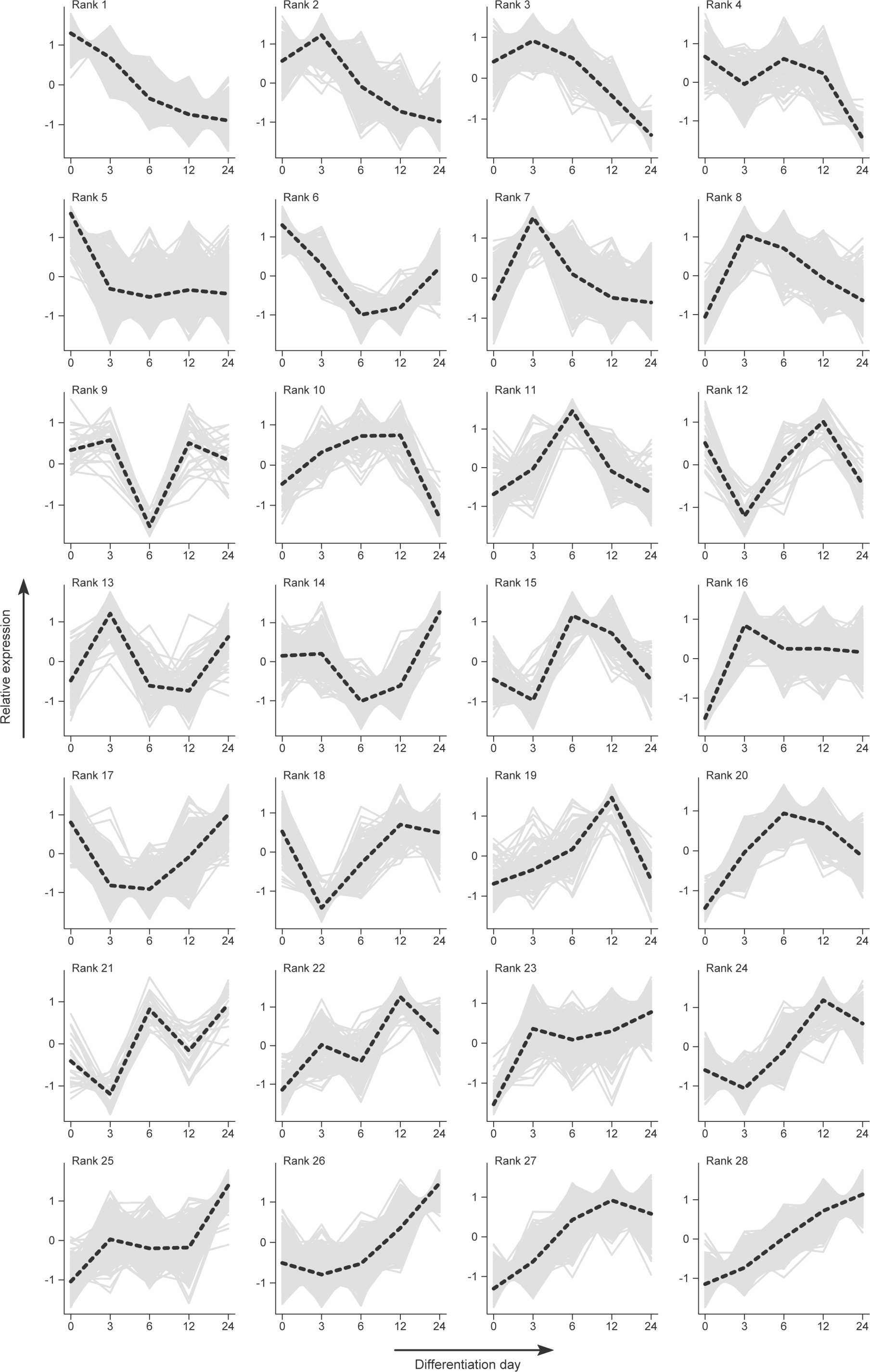
Ranking of gene expression trajectories in differentiating TRE-Ngn2 cells. Longitudinal RNA-seq data for control-treated TRE-Ngn2 cells were analyzed by DP_GP_cluster [53] and ranked from monotonic downregulation (rank 1) to monotonic upregulation (rank 28) based on the cluster-specific Kendall’s trend τ values. Solid gray lines, temporal dynamics of individual genes in each cluster. Dashed black lines, gene cluster trajectories.

**Figure S6.**
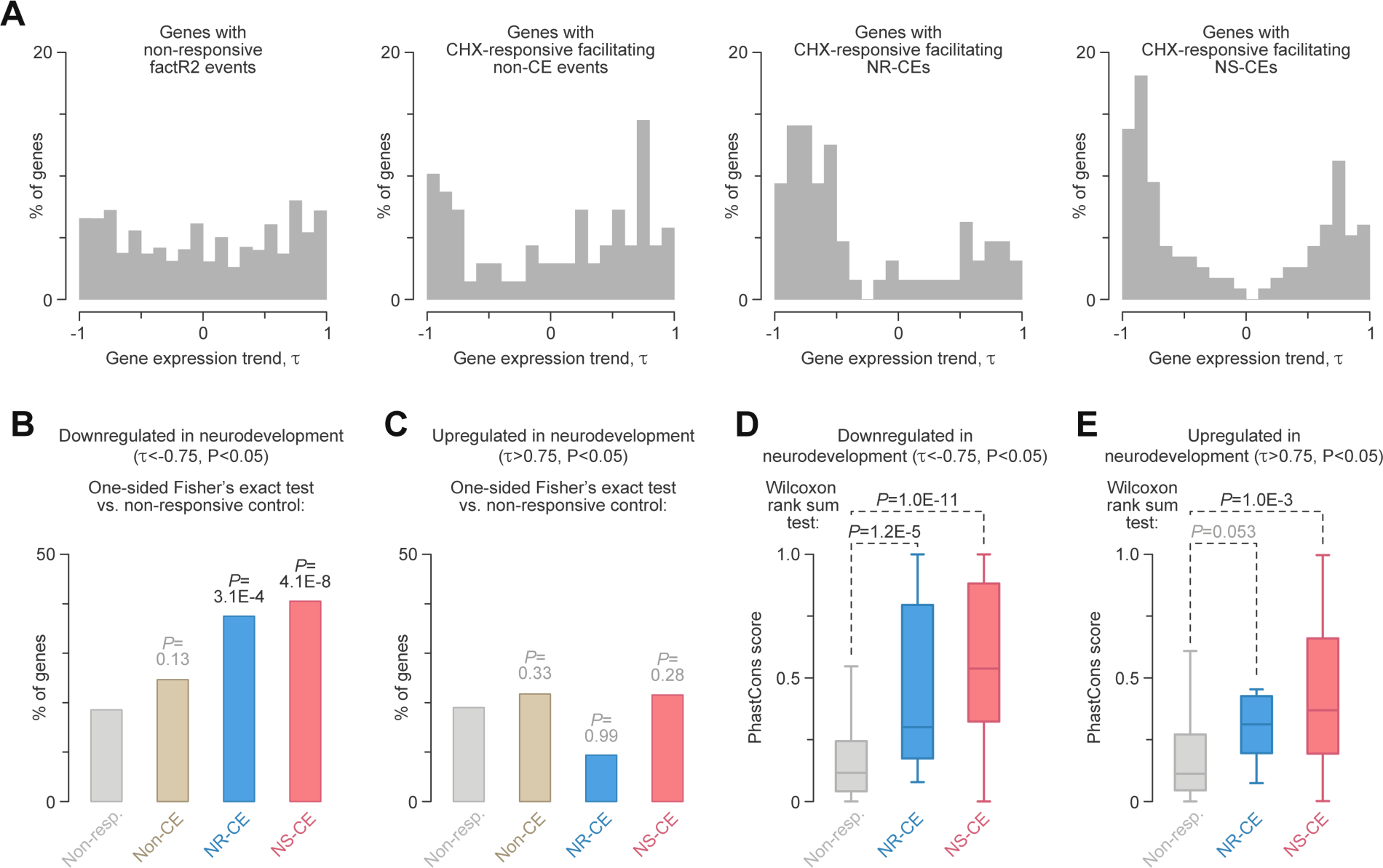
AS-NMD control of neurodevelopmentally downregulated genes. **(A)** Distribution of Kendall’s trend τ for individual genes with the indicated types of factR2 events. The prominent peaks of negative τ values in the [-1, -0.75] interval for genes with facilitating NR-CEs and NS-CEs suggest that these groups are often monotonically downregulated in developing neurons. **(B)** One-sided Fisher’s exact test showing that the monotonic downregulation trend (Kendall’s τ<-0.75, *P*<0.05) is significantly overrepresented among genes with CHX-responsive facilitating NR-CEs and NS-CEs compared to genes containing only non-responsive factR2 events. **(C)** No significant difference is detected for the monotonic upregulation trend (Kendall’s τ>0.75, *P*<0.05). **(D-E)** PhastCons [94] analysis of the four groups of events in (A-C) shows strong conservation of the intronic sequence context in NS-CEs in general and NS-CEs residing in monotonically downregulated genes in particular.

**Figure S7.**
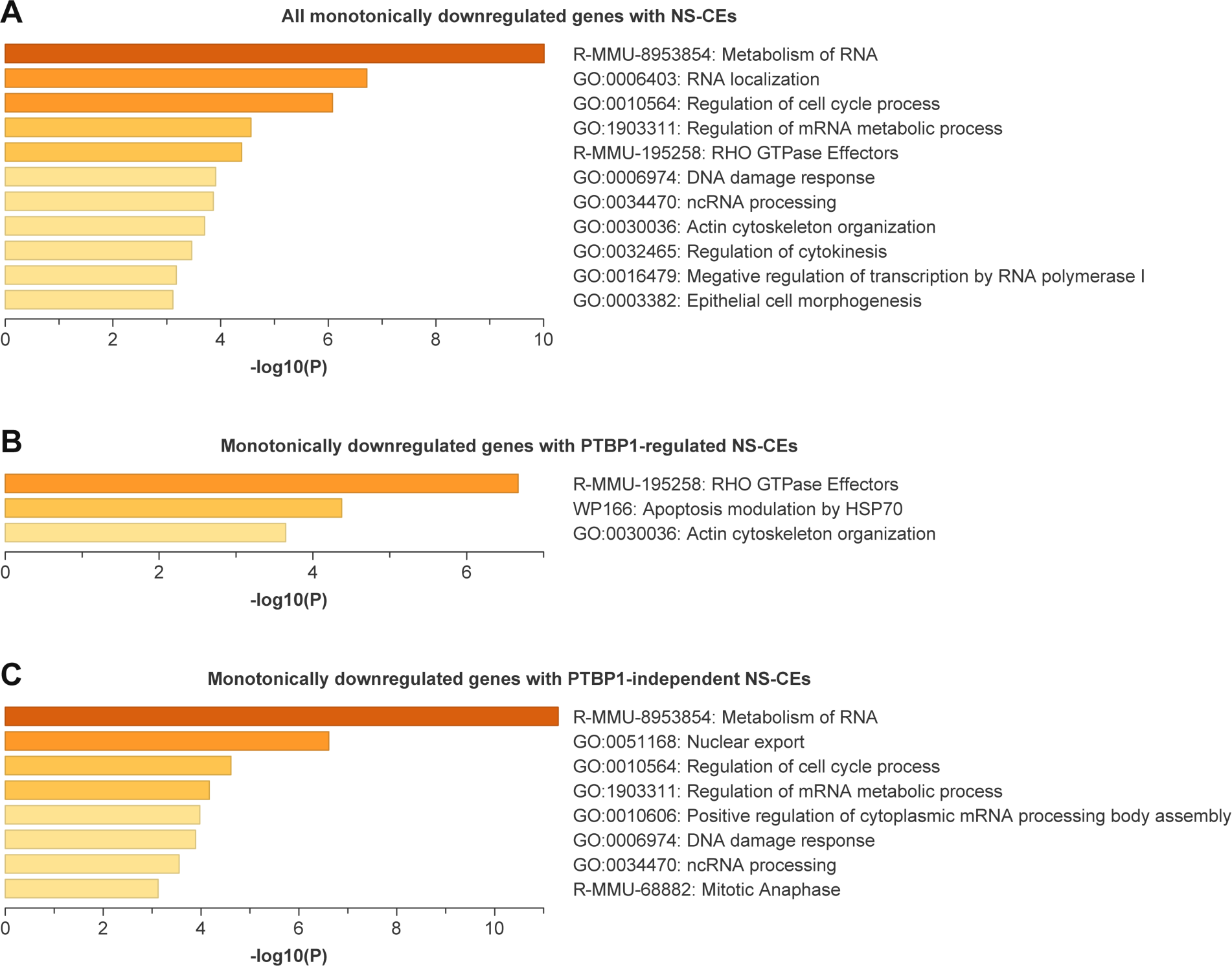
Neurodevelopmentally downregulated genes with facilitating NS-CEs are enriched for specific biological functions. Shown are significantly enriched (*P*<0.001) functional group summaries identified by Metascape [54] for **(A)** all monotonically downregulated genes (Kendall’s τ>0.75, *P*<0.05) containing NS-CEs; **(B)** monotonically downregulated genes with PTBP1-regulated NS-CEs; and **(C)** monotonically downregulated genes with PTBP1-independent NS-CEs. See Table S4 for further details.

**Figure S8.**
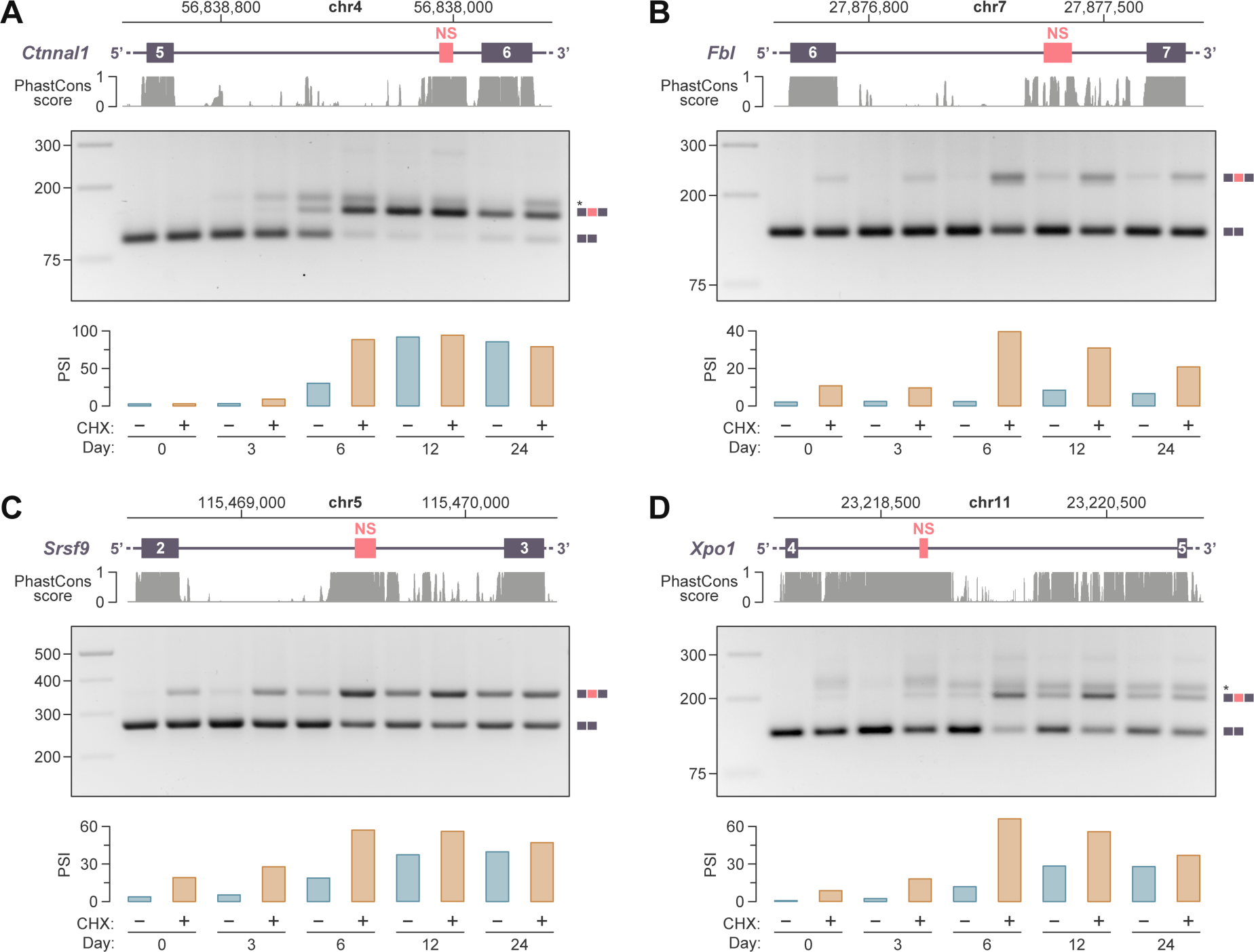
Experimental validation of facilitating NS-CEs encoded in genes monotonically downregulated in developing neurons. TRE-Ngn2 cultures were briefly treated with DMSO (odd samples) or CHX (even samples) on differentiation days 0-24 and analyzed by RT-PCR with primers flanking NS-CEs in (A) *Ctnnal1*, (B) *Fbl*, (C) *Srsf9*, and (D) *Xpo1* genes. In each panel, the NS-CE containing parts of genes and PhastCons vertebrate conservation tracks are shown on the top; gel analyses of the RT-PCR products are in the middle; and quantifications of NS-CE percent spliced in (PSI) values in DMSO (blue) and CHX (orange) treated samples are at the bottom. Note that in all four cases, NS-CE inclusion tends to increase as a function of development.

**Figure S9.**
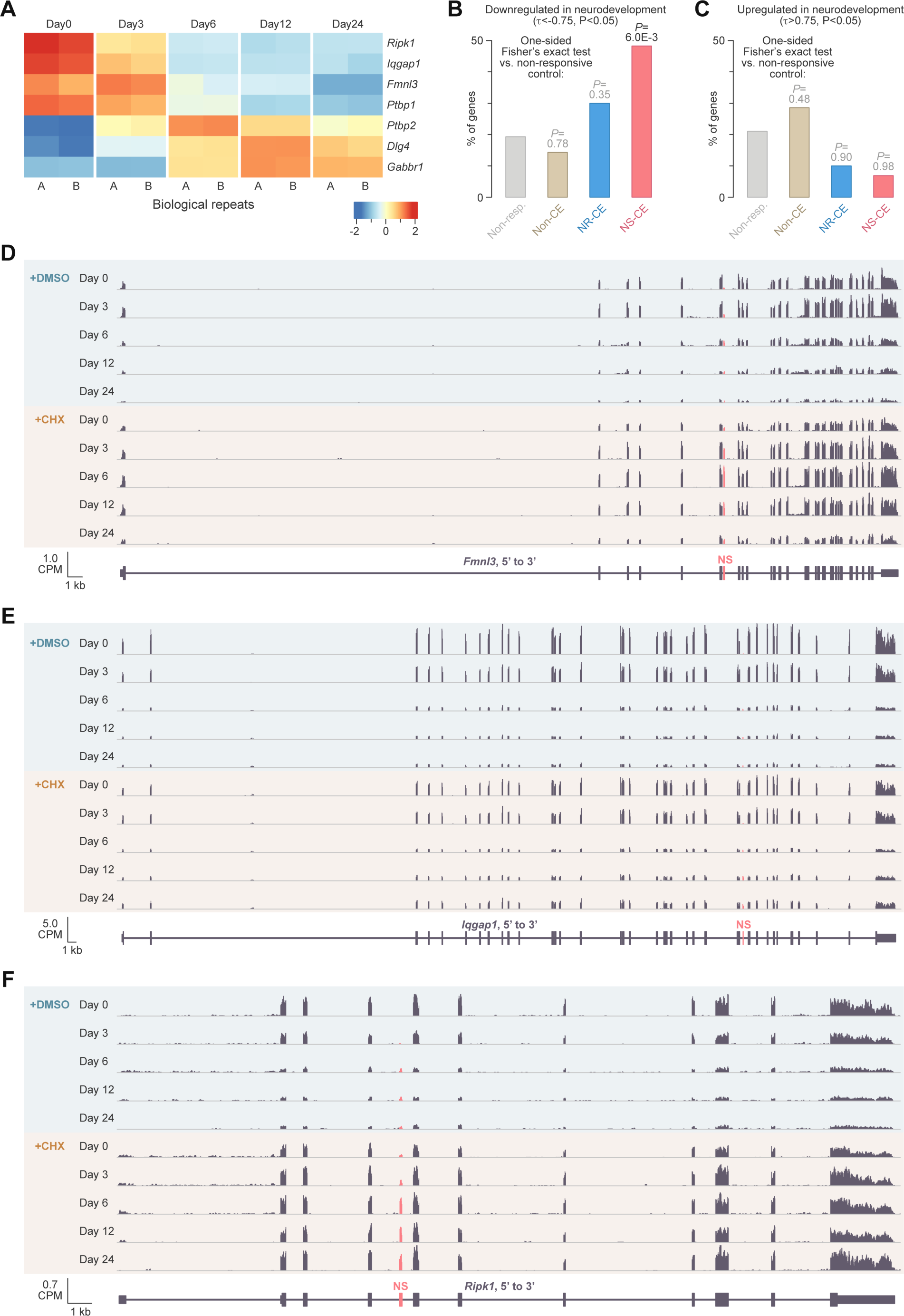
Examples of PTBP1-regulated AS-NMD target genes. **(A)** The expression dynamics of *Ptbp1* and PTBP1-cotrolled targets in differentiating TRE-Ngn2 cells. The targets shown include neurodevelopmentally upregulated genes with previously characterized NR-CEs (*Ptbp2*, *Dlg4*, and *Gabbr1*) and neurodevelopmentally downregulated genes with NS-CEs shortlisted in this study (*Fmnl3*, *Iqgap1*, and *Ripk1*) (Table S3). **(B)** One-tailed Fisher’s exact test showing that the monotonic downregulation trend (Kendall’s τ<-0.75, *P*<0.05) is overrepresented among genes with PTBP1-controlled facilitating NS-CEs compared to genes containing only non-responsive factR2 events. **(C)** Conversely, monotonic upregulation (Kendall’s τ>0.75, *P*<0.05) is not enriched among genes with different types of PTBP1-controlled AS-NMD events. **(D-F)** Count-per-million (CPM) normalized RNA-seq coverage plots for (D) *Fnml3*, (E) *Iqgap1*, and (F) *Ripk1*. The NS-CEs identified by factR2 are highlighted in red. All three genes are strongly downregulated in control-treated differentiating TRE-Ngn2 samples, and this effect is partially rescued by CHX.

**Figure S10.**
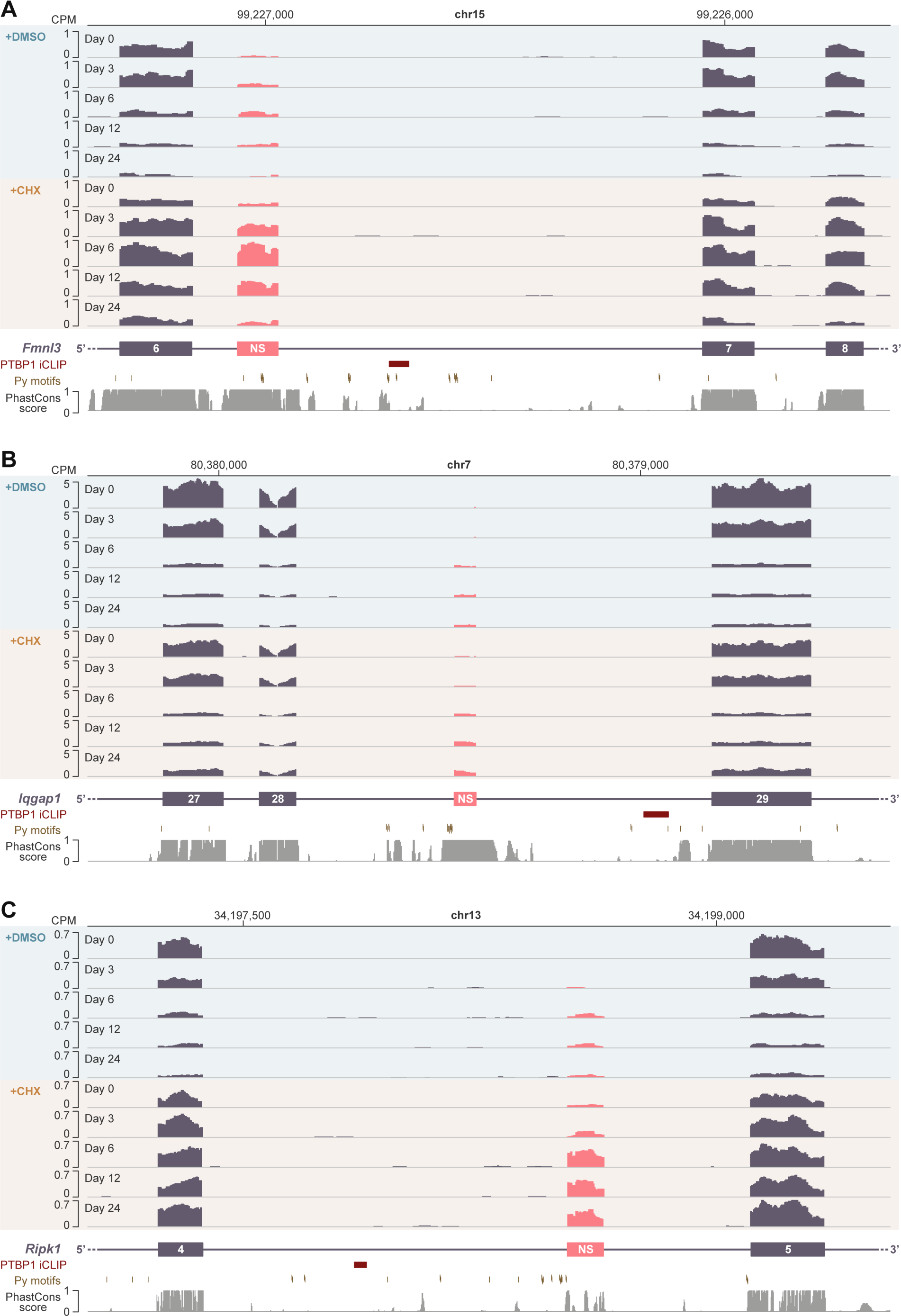
Alternatively spliced parts of neurodevelopmentally downregulated PTBP1 targets. Shown are CPM-normalized RNA-seq coverage plots for the regulated parts of **(A)** *Fnml3*, **(B)** *Iqgap1* and **(C)** *Ripk1*. The NS-CEs are highlighted in red. PTBP1 iCLIP clusters [55], PTBP1-specific motifs (Py; YTCTYY and YYTCTY), and vertebrate PhastCons tracks are shown at the bottom of each panel. The inclusion of NS-CEs increases as a function of development, and this effect is especially prominent in the CXH-treated series.

**Figure S11.**
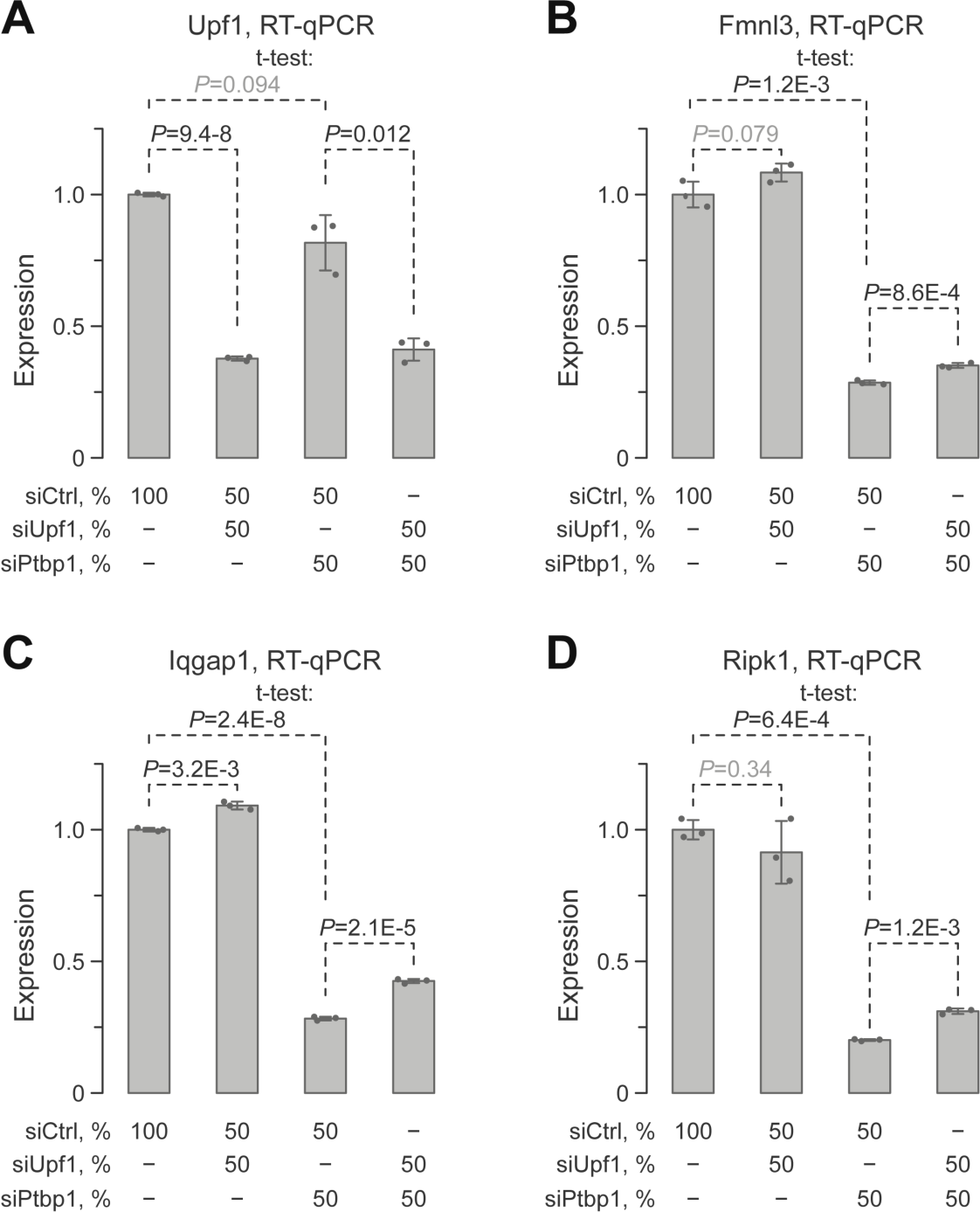
PTBP1 downregulation dampens the expression of *Fmnl3*, *Iqgap1* and *Ripk1* in an NMD-dependent manner. Mouse ESCs were treated for 48 hours with siRNAs indicated at the bottom of each panel and analyzed by RT-qPCR with primers against constitutively spliced parts of **(A)** *Upf1*, **(B)** *Fmnl3*, **(C)** *Iqgap1*, and **(D)** *Ripk1*. Note that PTBP1 knockdown dampens the expression of its target genes and that the knockdown of the key NMD factor UPF1 partially rescues this effect.

**Figure S12.**
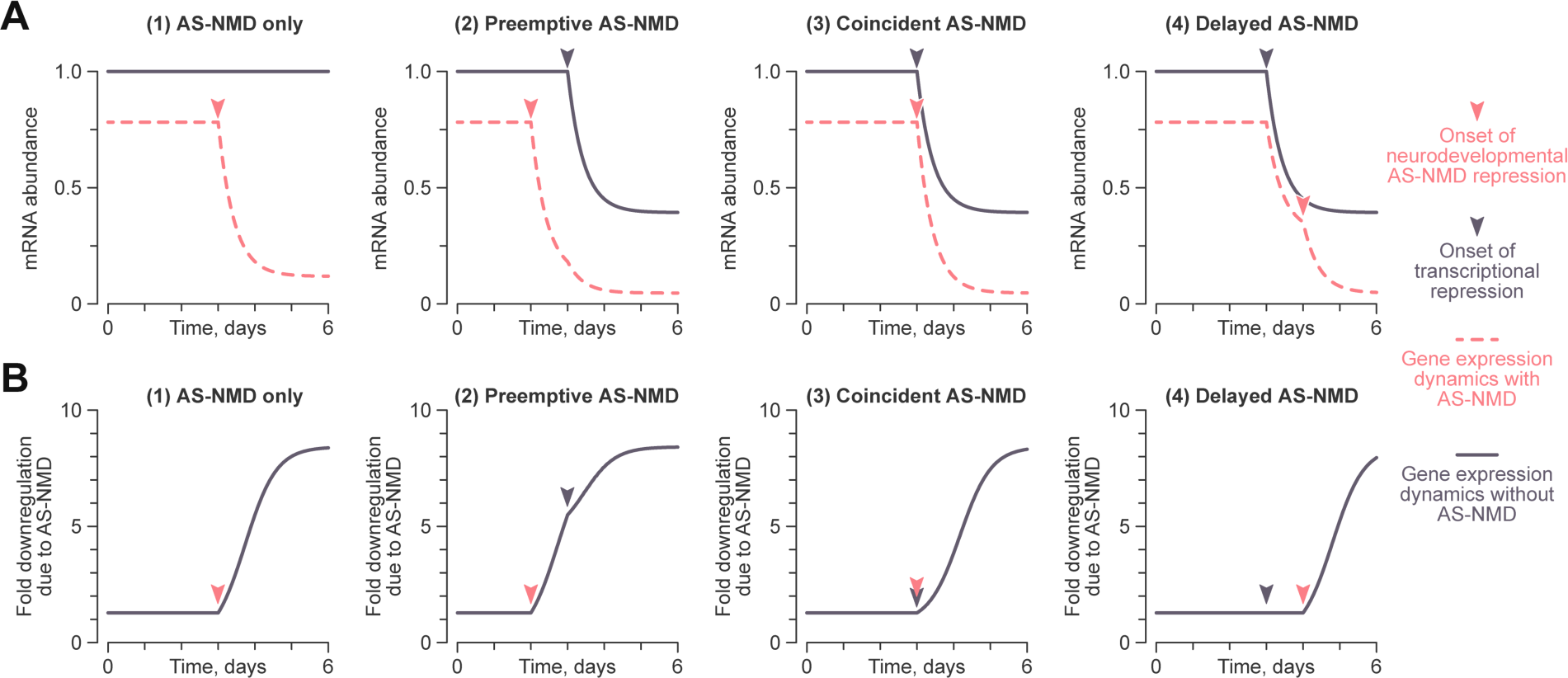
Possible effects of AS-NMD on gene downregulation in developing neurons. We used the ordinary differential equation model described in Materials and Methods to estimate AS-NMD contribution to gene expression dynamics when it (1) is the only downregulation mechanism or (2-4) functions alongside transcriptional repression with different onset times. **(A)** Gene expression dynamics with (red) and without (mauve) AS-NMD for the four regulation possibilities. **(B)** Effect of AS-NMD on gene expression calculated from (A) by normalizing the without-AS-NMD trajectories by their with-AS-NMD counterparts.

**Figure S13.**
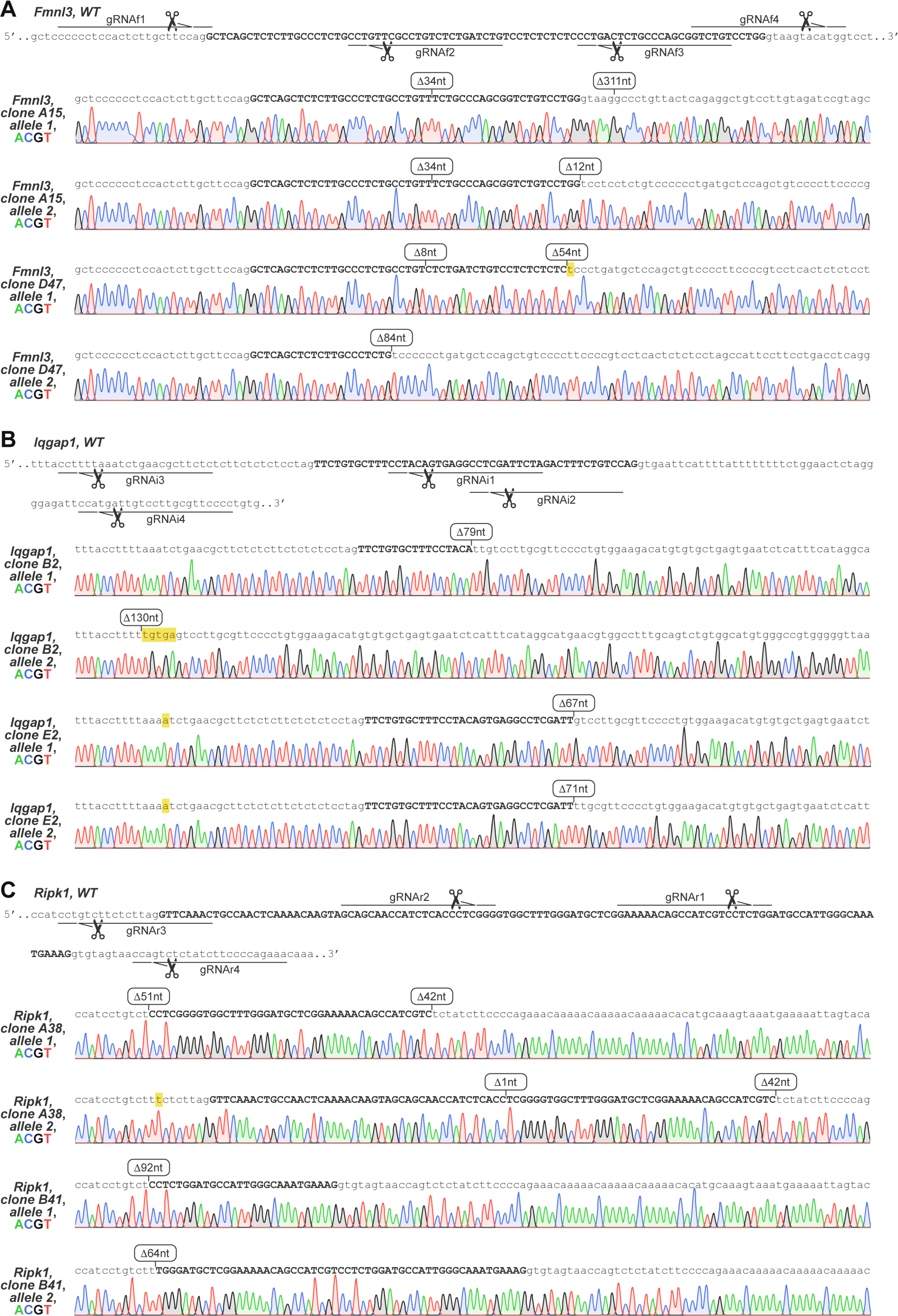
CRISPR/Cas9 disruption of PTBP1-regulated NS-CEs. TRE-Ngn2 ESCs were transfected with the expression constructs encoding Cas9 and four different CRISPR gRNAs targeting the NS-CEs and their adjacent intronic sequences in **(A)** *Fmnl3*, **(B)** *Iqgap1*, and **(C)** *Ripk1*. Two clones with biallelic mutations in the NS-CE region were selected for each gene and analyzed by Sanger sequencing. Each panel shows the wild-type gene sequence and the gRNA positions at the top and allele-specific sequencing chromatograms for the two mutant clones at the bottom. Upper case, wild-type NS-CEs and their remnants. Lower case, intronic sequences. Rounded rectangles, CRISPR/Cas9-induced deletions. Yellow highlight, insertions and nucleotide substitutions.

**Figure S14.**
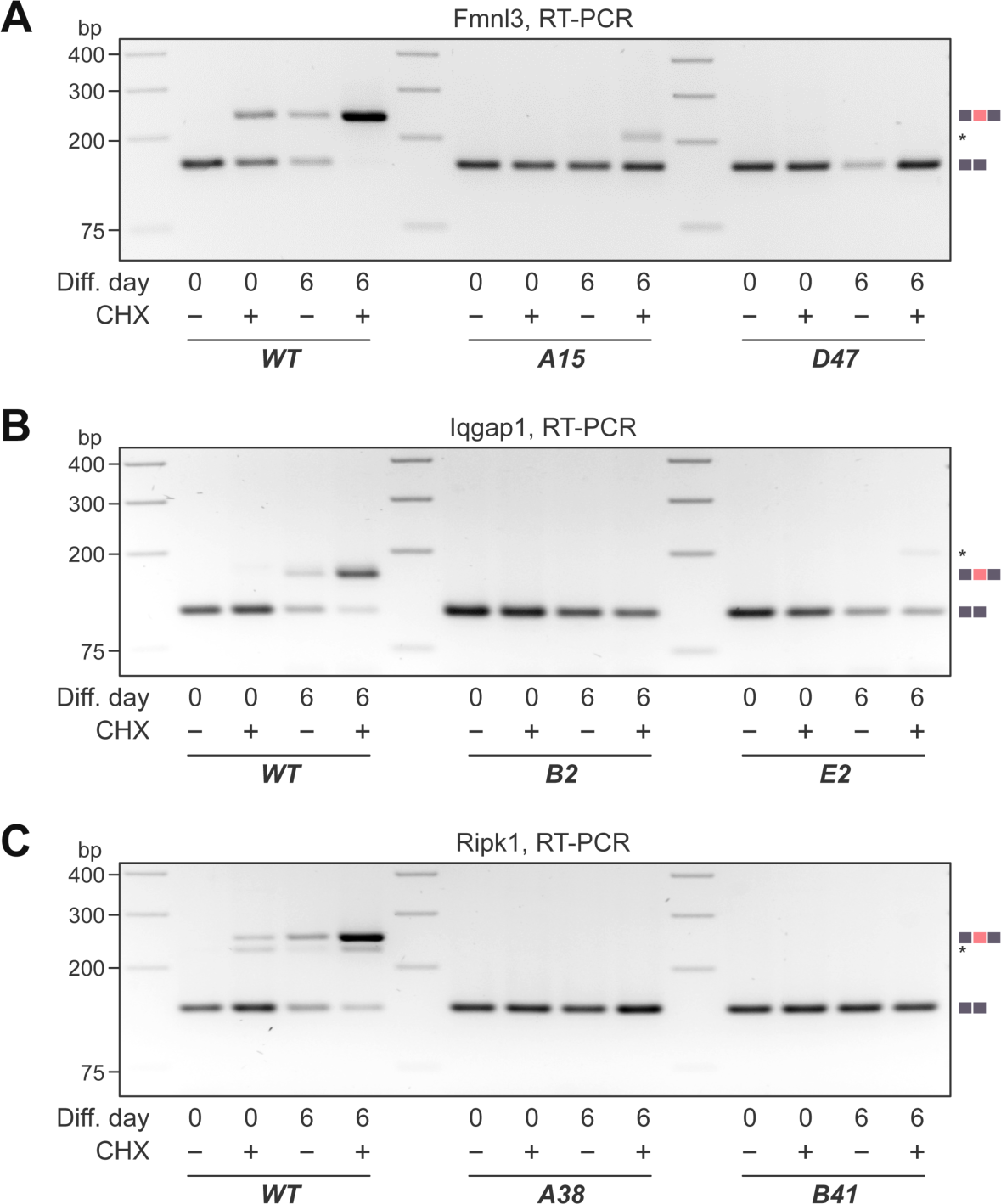
RT-PCR analysis of wild-type and mutant TRE-Ngn2 cells. Wild-type (WT) TRE-Ngn2 cells and the NS-CE mutants introduced in Fig. S13 were treated with DMSO or CHX on differentiation days 0 (ESCs) or 6 (young neurons) and analyzed by RT-PCR with **(A)** *Fmnl3*-, **(B)** *Iqgap1*- or **(C)** *Ripk1*-specific primers designed against the NS-CE-flanking exons. In the wild type, NS-CE-containing mRNAs are readily detectable on day 6 in DMSO-treated samples, becoming a major splice form in the presence of CHX. The ability to produce NMD-sensitive mRNA isoforms is either completely lost (clones D47, B2, A38 and B41) or drastically impaired (clones A15 and E2) in the mutant cells. Asterisks, unspecific amplification products and low-abundance NMD-sensitive transcripts detectable in clones A15 and E2.

